# Upregulation of ATP-purinergic P2x2 receptors in the cochlea over-amplifies hearing sensitivity leading to hyperacusis and attenuation by antagonists

**DOI:** 10.64898/2026.06.17.733049

**Authors:** Tian-Ying Zhai, Chun Liang, Jin Chen, Jie Yang, Yong Kong, Yan Zhu, Ning Yu, Hong-Bo Zhao

## Abstract

Hearing hypersensitivity (hyperacusis) is a common hearing stress and can cause many psychological diseases, e.g., anxiety, learning disabilities, and attention-deficit/hyperactivity disorder (ADHD). Here, we report an unexpected finding that the upregulation of P2x2 ATP-purinergic receptors in the cochlea links to hyperacusis generation. We found that P2x2 expression in the cochlea but not in auditory centers was upregulated in the hyperacusis generated by Cx26 deficiency. Overexpression of P2x2 in the cochlea also caused hyperacusis. Conversely, downregulation of P2x2 expression or administration of P2x2 antagonists attenuated hyperacusis. We further found that upregulation of P2x2 receptors in the cochlea increased outer hair cell (OHC) electromotility through the post-transcription functional modulation to potentiate active cochlear amplification leading to hearing hypersensitivity. Such enhancements in OHC electromotility and active cochlear amplification were also suppressed by P2x2 receptor antagonists. Overall, these findings demonstrate that P2x2-mediated ATP-purinergic signaling in the cochlea plays a critical role in hyperacusis generation; targeting P2x2 receptors can attenuate hyperacusis stress, which may also offer a therapeutic strategy for other related psychological comorbidities.

**Significance statement:** Hearing hypersensitivity is a common hearing stress and can cause many other psychological disorders. However, little is known about the underlying genetic and cellular mechanisms. Also, it lacks efficient drugs for their treatments in the clinic. In this study, we found that upregulation of P2x2 ATP-purinergic receptors in the cochlea can potentiate outer hair cell electromotility, which is an active cochlear amplifier in mammals and can increase hearing sensitivity and frequency selectivity, through post-transcription functional modulation to enhance active cochlear amplification leading to hearing hypersensitivity. These enhancements can be inhibited by administrations of P2x2 receptor antagonists both *in vitro* and *in vivo*. These findings revealed a new genetic and cellular mechanism underlying hyperacusis generation and opened a new avenue to develop an efficient therapy for this common hearing stress and other associated psychological comorbidities.

## Introduction

Hyperacusis (hearing hypersensitivity) is a common hearing stress and can cause many psychological disorders, such as anxiety, learning disabilities, attention-deficit/hyperactivity disorder (ADHD), and post-traumatic stress disorder (Baguley, 2003; Pienkowski et al., 2014; Tyler et al., 2014; Xie et al., 2025). Hyperacusis is also associated with other hearing dysfunctions, such as tinnitus, loudness recruitment, and age-related hearing loss (Knipper et al., 2013; Xie et al., 2025). However, despite its prevalence, little is known about the mechanisms underlying hyperacusis generation; no clear genetic cues have been identified, and no specific, effective pharmacological treatments are available in the clinic.

ATP-purinergic cell signaling is a signaling pathway that extracellular ATP activates purinergic (P2) receptors on the cell surface to influence cellular functions and has a vital role in a wide array of physiological and pathological processes, such as pain, inflammation, neurotransmission, and so on (North, 2002; Surprenant & North, 2009). P2 receptors have two subgroups: ATP-gated ionotropic (P2x) and G protein-coupled metabotropic (P2y) subgroups. The P2x receptor has 7 subtypes (P2x1-7), which are widely expressed in various tissues and cells functioning in many cellular, physiological, and pathological processes (North, 2002; Surprenant & North, 2009). ATP-purinergic signaling also plays an important role in hearing function. In earlier studies, it was found that ATP can elevate intracellular Ca^++^ concentration in hair cells to modify sound transduction and neurotransmission (Ashmore & Ohmori, 1990; Dulon et al., 1991; Housley et al., 1999; Sugasawa et al., 1996). Later, it was found that ATP can mediate hearing sensitivity, extend the dynamic range of hearing, synchronize auditory nerve activity in development, mediate type II auditory nerve activity, and contribute to hair cell damage response (Housley et al., 2013; Liang et al., 2025; Liu et al., 2015; Telang et al., 2010; Thorne et al., 2004; Tritsch & Bergles, 2010; Tritsch et al., 2007; Weisz et al., 2009). We also found that ATP-purinergic signaling can mediate outer hair cell (OHC) electromotility, gap junctional coupling, K^+^-recycling, and endocochlear potential (EP) generation, and cochlear efferent suppression (Chen et al., 2015; Liang et al., 2025; Yu & Zhao, 2008; Zhao et al., 2005; Zhu & Zhao, 2010, 2012). The fact that P2x2 mutations can cause nonsyndromic hearing loss DFNA41 (Faletra et al., 2014; Yan et al., 2013; Zhu et al., 2017) further indicated that ATP-purinergic signaling has a critical role in hearing function.

In this study, we report an unexpected finding that the upregulation of P2x2 expression is associated with hyperacusis generation. Overexpression of P2x2 in the cochlea produced hyperacusis through increasing outer hair cell electromotility to enhance active cochlear amplification over-amplifying hearing sensitivity, whereas downregulation of P2x2 expression or administration of P2x2 antagonist could attenuate enhancement and hyperacusis. These findings revealed that P2x2-mediated ATP purinergic cell signaling has a critical role in hyperacusis generation; targeting P2x2 receptors can overcome hyperacusis.

## Results

### Upregulation of P2x2 receptors in the cochlea in hyperacusis generated by Cx26 deficiency

Fig. 1 shows the upregulation of P2x2 receptors in the cochlea in hyperacusis generated by Cx26 hetero-deletion (Cx26^+/-^). Fig. 1**a-f** shows hyperacusis generation in Cx26^+/-^ mice. Acoustic startle response (ASR) behavior test in Cx26^+/-^ mice demonstrated an enhanced response (Fig. 1**a-c**). The amplitude of ASR was significantly increased in comparison with WT mice (Fig. 1**b&c**). Also, as described before (Liu et al., 2023), ABR thresholds in Cx26^+/-^ mice were significantly reduced, and CM and DPOAE in Cx26^+/-^mice were significantly increased in comparison with those in WT mice (Fig. 1**d-f**). In RNA-Seq examination (Fig. 1**g&h**), it was found that expression of P2x2 receptors in the cochlea in Cx26^+/-^ mice was significantly upregulated. The increase was 0.3989±0.077 log2Fold upregulation (Fig. 1**h**). Droplet digital PCR (dPCR) further confirmed that P2x2 expression in the Cx26^+/-^ cochlea had 2.37±0.19 fold-increase (P<0.001, t test, 2-tailed) in comparison with WT mice (an inset in Fig. 1**g**). Immunofluorescent staining for P2x2 (Fig. 1**i**) showed strong P2x2 labeling in the organ of Corti (OC), spiral limbs (SLM), outer sulcus cells (OSCs), and the cochlear lateral wall. Especially, apparent P2x2 labeling is visible at the outer hair cell (OHC) and inner hair cell (IHC).

**Fig. 1.**
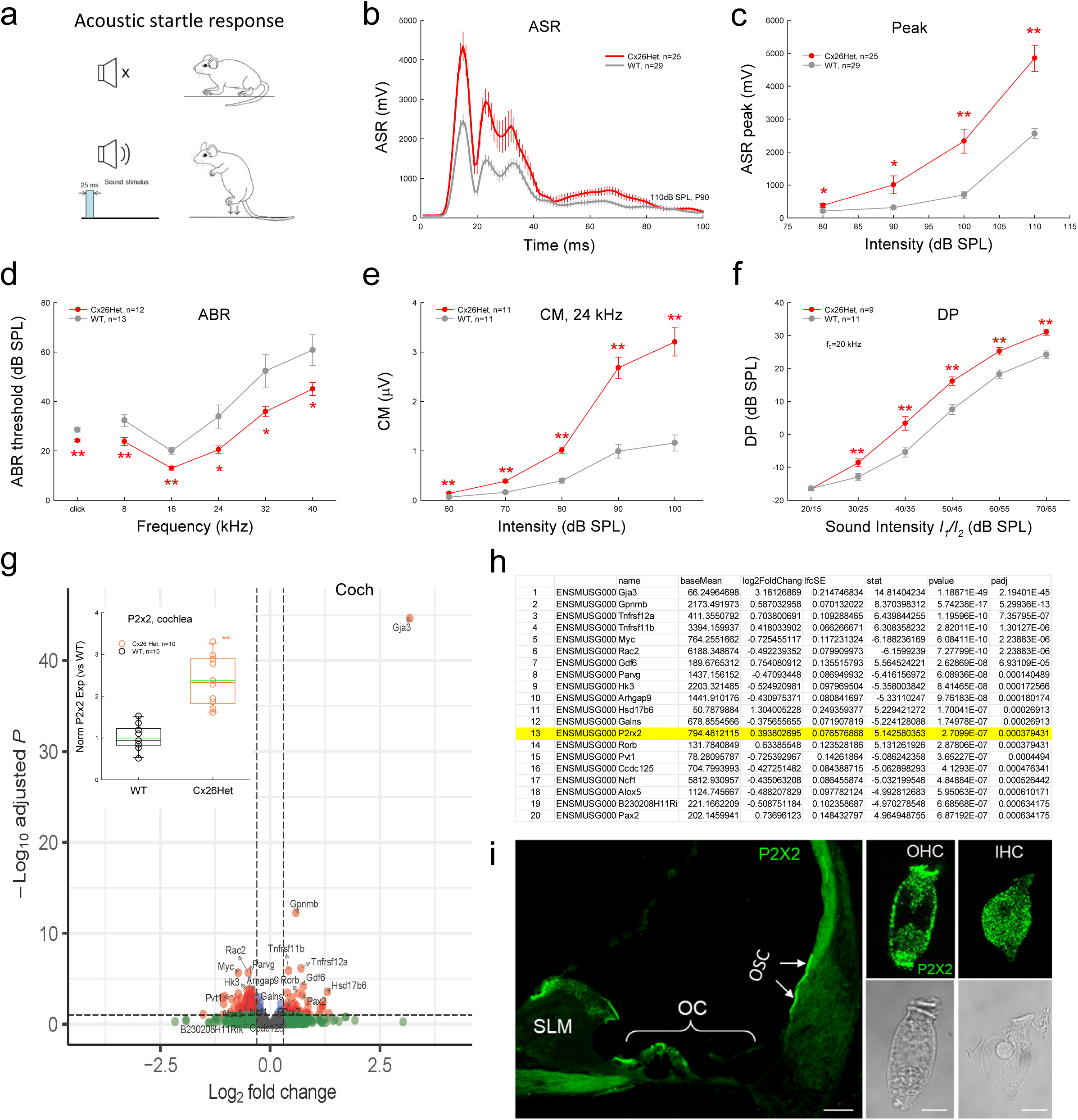
Upregulation of P2x2 expression in the cochlea in hyperacusis generated by Cx26 heterozygous deletion. **a-c**: Enhanced acoustic startle response (ASR) in Cx26 heterozygous deletion (Cx26^+/-^) mice. Panel **a**: diagram of ASR measurement. Panel **b**: the averaged ASR trace evoked by 110 dB noise pulse. **d-f**: Hearing over-sensitivity in Cx26^+/-^ mice measured by ABR threshold, CM, and DPOAE recordings. In Cx26^+/-^ mice, ABR thresholds were significantly reduced; CM and DPOAE were significantly increased. **g**: Volcano plot of gene upregulation and downregulation in the cochlea in Cx26^+/-^ mice in Bulk Poly(A) RNA-Seq examination. P2x2 expression in the cochlea in Cx26^+/-^ mice was significantly upregulated. Insect: Upregulation of P2x2 expression in the cochlea in Cx26^+/-^ mice assessed by digital droplet PCR (dPCR). P2x2 expression in the cochlea in Cx26^+/-^ mice was normalized to that in WT mice. **h**: Changes of top 20 genes ranked by p-values in the cochlea in Cx26^+/-^ mice in RNA-Seq examination. **i**: Expression of P2x2 in the cochlea examined by immunofluorescent staining. Apparent P2x2 labeling is visible at inner hair cells (IHCs) and outer hair cells (OHCs) in the organ of Corti (OC), spiral limbs (SLM), and outer sulcus cells (OSCs) in the lateral wall. Scale bars: 100 µm and 10 µm. *: P<0.05, **: P<0.01, 2-tail t test.

However, there were neither expressions nor upregulations of P2x2 receptors in auditory centers in Cx26^+/-^ mice (Fig. S1-2). The expressions of P2x2 in the auditory cortex (AC), inferior colliculus (IC), and cochlear nucleus (CN) in Cx26^+/-^ mice were not detectable at the transcriptional level by dPCR examination (Fig. S2). P2x2 expressions in the AC, IC, and CN in Cx26^+/-^ mice were also not significantly altered in RNA-Seq examination (Fig. S1). P2x7 has expression in the cochlea and deficiency of P2x7 can induce hearing hypersensitivity (Liang et al., 2025). P2x7 expression in the AC, IC, CN and cochlea also had no significant changes in Cx26^+/-^ mice (Fig. S3).

### Enhanced hearing sensitivity induced by overexpression of P2x2 in the cochlea

We further tested whether upregulation of P2x2 expression in the cochlea in the WT mice could increase hearing sensitivity (Fig. 2). The right cochlea of the WT mice was injected with AAV-P2x2 vectors via the posterior semicircular canal (PTSC) to increase P2x2 expression in the cochlea, and the left cochlea was not injected serving as an internal control. AAV empty vector without P2x2 was also used for injections as control. After injection of AAV-P2x2 vectors, the expression of P2x2 in the right injection cochlea had 2.75±0.39 fold significantly increasing (Fig. 2**c**, P=0.0017, one-way ANOVA with a Bonferroni correction), similar to P2x2 upregulation (2.27±0.17 fold-increase) in Cx26^+/-^ mice (the inset in Fig. 1**g**). Also, the expression pattern is similar to the native expression of P2x2 in the cochlea (Fig. 1**i**); the injected AAV-P2x2 vectors had intensive expression at IHCs and OHCs (Fig. 2**a&b**). Especially, the expression of P2x2 at the OHC lateral wall showed a ring labeling structure (Fig. 2**a**). Auditory function testing shows that hearing sensitivity in the right ear with AAV-P2x2 vector injection was significantly increased; ABR thresholds were reduced (Fig. 2**d**) and CM and DPOAE were increased (Fig. 2**e&f**). Behavioral tests measured by ASR also showed that hearing sensitivity was increased (Fig. 2**g&h**). The increase demonstrated dose-dependence (Fig. 2**i**). The EC_50_ concentration was 5.6e+12 vg/mL (Fig. 2**i**). However, injection of empty AAV vector without P2x2 had no effect on hearing function, and the ASR behavioral testing had no significant changes in comparison with the control group without any surgery injection (Fig. 2**d-h**). The expression of P2x2 in the cochlea with empty AAV vector injection also had no significant change (Fig. 2**c**).

**Fig. 2.**
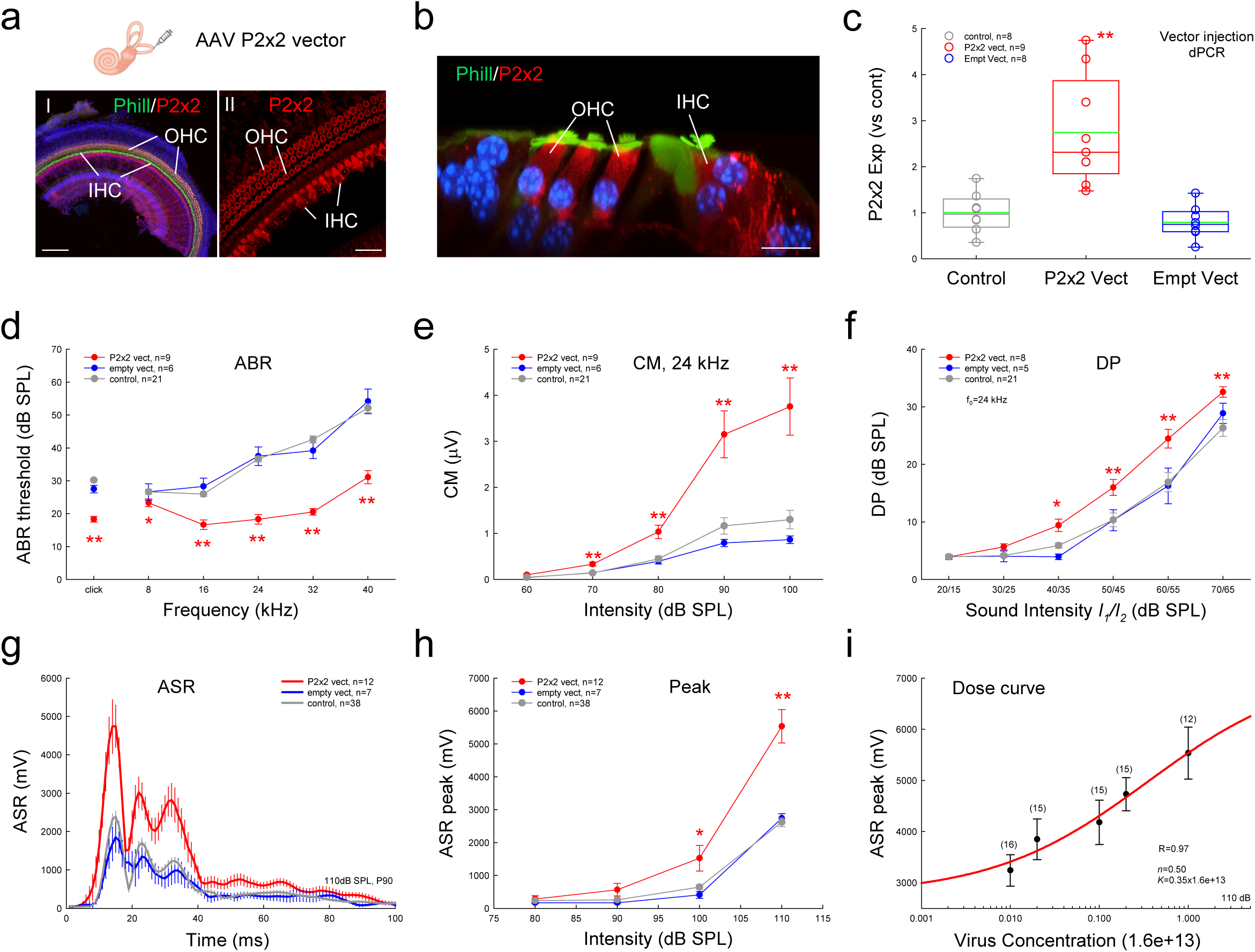
Generation of hyperacusis by overexpression of P2x2 in the cochlea. AAV vectors were micro-injected into the cochlea via the posterior semicircular canal (PTSC). Only right ear was injected. **a-b**: Expression of P2x2 in the cochlea after injection of AAV P2x2 vectors into the cochlea. The expression of P2x2 is mainly located at IHCs and OHCs. Insect: diagram of injection of AAV vectors into inner ear via the PTSC. **c**: The expression of P2x2 in the cochlea after injection of AAV P2x2 vectors or AAV empty vectors measured by dPCR. The P2x2 expressions were normalized to control mice without injection. Injection of P2x2 vectors but not empty vectors significantly increased P2x2 expression in the cochlea. **d-f**: Hearing function in the right ear measured by ABR, CM, and DPOAE with the close-field recording. Hearing sensitivity in P2x2 vector injected ears is significantly increased by reduced ABR thresholds and increased responses in CM and DPOAE. **g-h**: Increase of ASR in P2x2 vector injected mice but not empty vector injected mice. **i**: Dose curve of ASR increases for P2x2 vector injection. Numbers within parentheses at each point represent the measured animal numbers. A red smooth line represents amplitudes of ASR peaks fitting to a Hill’s function: ASR = a * *C^n^*/ (*K^n^* + *C^n^*) + y0, where *n*=0.50, *K*=0.35 (x1.6e+13) i.e., EC_50_=0.35 x 1.6e+13=5.6e+12 vg/mL. *: P<0.05, **: P<0.01, one-way ANOVA with a Bonferroni correction in panel **c**, and 2-tail t test in panel **d-h**.

We also examined Cx26 and P2x7 expressions after injection of AAV-P2x2 vectors. After injection of AAV-P2x2 vectors, Cx26 and P2x7 expressions in the right cochlea had no significant changes in comparison with the left ear without any injection (Fig. S4, P=0.75-0.81 one-way ANOVA). Also, referring to the left ears without injection, the expression of P2x2 in the right ears with injection of AAV-P2x2 vectors was significantly increased by 2.85±0.43 folds (Fig. S4, P=0.0028, one-way ANOVA with a Bonferroni correction), similar to 2.75±0.39 folds of the increment referring to the control group without any injection (Fig. 2**c**).

### Elimination of hyperacusis by down-regulation of P2x2 expression

To further investigate the role of P2x2-mediated cell signaling pathway in hyperacusis generation, we tested whether downregulation of P2x2 receptors can eliminate the hyperacusis generation (Fig. 3). We crossed Cx26^+/-^ mice with P2x2^+/-^heterozygous KO mice to downregulate P2x2 expression in the Cx26^+/-^ mice (Fig. 3**a**). After breeding with P2x2^+/-^ mice, the expressions of P2x2 in the cochlea in Cx26^+/-^ mice, P2x2^+/-^/Cx26^+/-^ mice, and P2x2^+/-^ mice were 2.05±0.29, 0.76±0.03, and 0.58±0.06 folds, respectively, referring to WT mice (Fig. 3**b**). There was no significant difference between P2x2^+/-^/Cx26^+/-^ mice and WT mice (P=0.16, one-way ANOVA). Along with the reduction of P2x2 expression in P2x2^+/-^/Cx26^+/-^ mice, the hearing function and ASR behavioral test in the P2x2^+/-^/Cx26^+/-^ mice returned to normal WT levels (Fig. 3**c-h**). Interestingly, the expression of P2x2 in the cochlea in P2x2^+/-^ mice (Fig. 3**b**) was significantly reduced to the half value (0.58±0.06 folds) referring to WT mice (P=0.03, one-way ANOVA with a Bonferroni correction). However, the hearing function and ASR behavioral testing in P2x2^+/-^ mice had no significant changes (Fig. 3**c-h**)

**Fig. 3.**
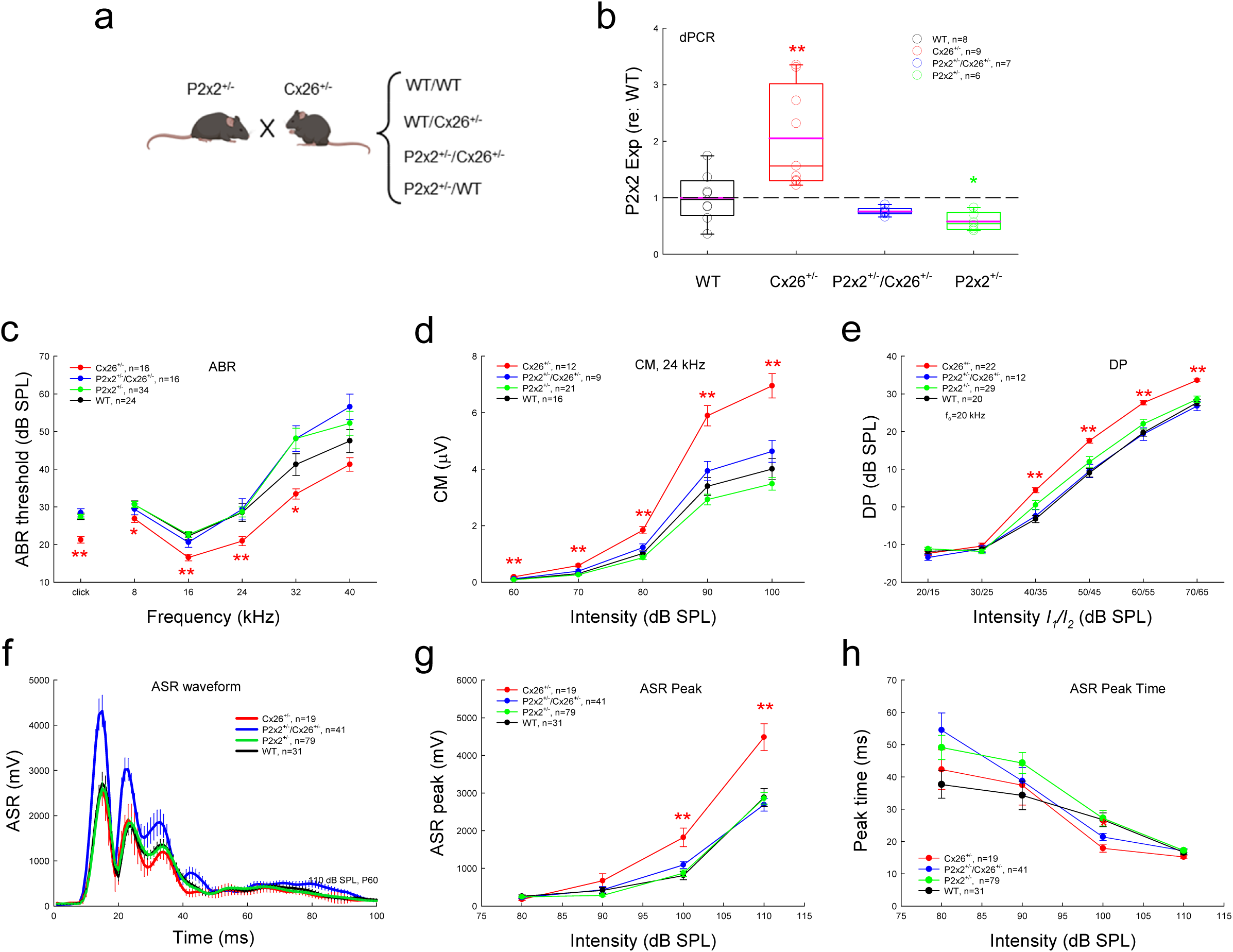
Elimination of hearing hypersensitivity in Cx26^+/-^ mice by down-regulation of P2x2 expression with crossing P2x2 KO mice. **a**: Diagram of crossing Cx26^+/-^ mice with P2x2^+/-^ heterozygous KO mice to down-regulate P2x2 expression. There are four genotypic groups after crossing: P2x2^+/-^/Cx26^+/-^, P2x2^+/-^/WT (P2x2^+/-^), WT/Cx26^+/-^(Cx26^+/-^), and WT/WT (WT). **b**: P2x2 expressions in the cochlea examined by dPCR. The levels of P2x2 expressions were normalized to WT mice. There is neither an increase of P2x2 expression in the cochlea in P2x2^+/-^/Cx26^+/-^ mice nor a significant difference between P2x2^+/-^/Cx26^+/-^ mice and WT mice, although there is a significant increase of P2x2 expression in the cochlea in Cx26^+/-^ mice. As expected, the expression of P2x2 in the P2x2^+/-^ mouse cochlea is 0.58±0.06 and significantly reduced to a half value referring to WT mice. **c-e**: Hearing hypersensitivity is eliminated after downregulation of P2x2 in P2x2^+/-^/Cx26^+/-^ mice. Hearing functions were measured by ABR, CM, and DPOAE. Hearing functions in P2x2^+/-^ mice also have no significant changes. **f-g**: ASRs in P2x2^+/-^/Cx26^+/-^ mice are significantly reduced in comparison with Cx26^+/-^ mice but there are no significant differences in ASRs between P2x2^+/-^/Cx26^+/-^ mice and WT mice. Also, ASR in P2x2^+/-^ mice has no significant change. **h**: There are no significant differences in ASR peak time among 4 groups. *: P<0.05, **: P<0.01, one-way ANOVA with a Bonferroni correction.

### Attenuation of hyperacusis by administration of P2x2 antagonist in Cx26^+/-^ mice

Administration of P2x2 receptor antagonists also could attenuate hyperacusis generation in Cx26^+/-^ mice (Fig. 4). PPADS (pyridoxalphosphate-6-azophenyl-2’,4’-disulfonic acid) is a P2x2 receptor antagonist. Both Cx26^+/-^ mice and WT mice were intraperitoneally injected with PPADS (1 mM, 0.1 mL/10g). ASR and hearing function were recorded before and after 8 h of injection. ASR was recorded again at 72 h after injection to see the recovery, since anesthesia is not required for its recording. Fig. 4**a-f** show that the enhanced ASR in Cx26^+/-^ mice was significantly reduced after injection of PPADS (Fig. 4**a-c**), while ASR in WT mice had no significant changes for PPADS injection (Fig. 4**d-e**). At 8 h after PPADS injection, the ASR in Cx26^+/-^ mice was reduced to the WT mouse level and had no significant difference between them (P=0.18, t test, 2-tailed) (Fig. 4**c**). The reduction was reversible. At 72 h after injection, the ASR in Cx26^+/-^mice was recovered and had significant enhancement again as pre-injection referring to WT mice (Fig. 4**a-c**). However, there were no significant changes in ASR peak times between Cx26^+/-^ mice and WT mice before and after PPADS administration (P=0.24-0.78, t test, 2-tailed) (Fig. 4**f**).

**Fig. 4.**
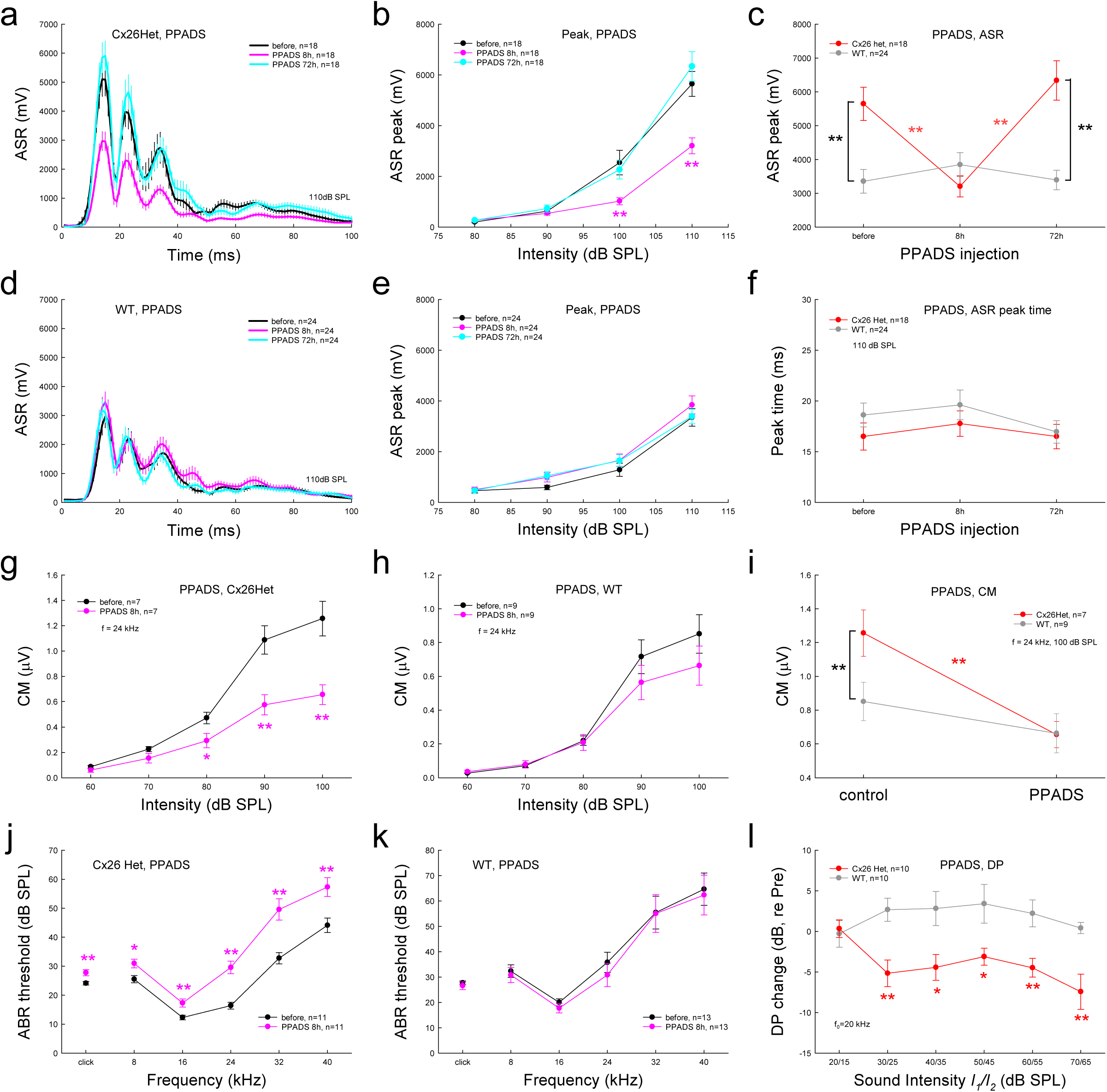
Elimination of hyperacusis in Cx26^+/-^ mice by administration of a P2x2 receptor antagonist PPADS. In Cx26^+/-^ mice, PPADS (1 mM) was intraperitonially injected into mice (0.1 mL/10g weight). ASR and hearing functions were measured before and after injection at 8 h. SAR was also measured again at post-injection 72 h to assess reversibility. **a-c**: Reversible elimination of increases of ASR in Cx26^+/-^ mice by administration of PPADS. The ASR was significantly reduced at 8 h after administration of PPADS and recovered after 72h of administration of PPADS. **d-e**: No significant effect of PPADS on ASR in WT mice. **f**: There is no significant effect of PPADS on the ASR peak time in Cx26^+/-^ mice and WT mice. **g-i**: Enhancement of CM in Cx26^+/-^ mice was eliminated after administration of PPADS. CM in Cx26^+/-^ mice was significantly reduced to the WT mouse level after administration of PPADS, while WT mice had no significant reduction after administration of PPADS (panel **i**). **j-k**: Elimination of hearing hypersensitivity in Cx26^+/-^ mice by PPADS. ABR thresholds were significantly increased in Cx26^+/-^ mice (panel **j**) but had no change in WT mice (panel **k**) after administration of PPADS. **l**: Significant reduction of DPOAE in Cx26^+/-^ mice after administration of PPADS. *: P<0.05, **: P<0.01, 2-tail t test.

Hearing functions in Cx26^+/-^ mice but not WT mice also show significant suppression after injection of PPADS (Fig. 4**g-l**). CM in Cx26^+/-^ mice but not in WT mice was significantly reduced after injection of PPADS (Fig. 4**g-i**). At 8 h after the injection of PPADS, the CM in Cx26^+/-^ mice was significantly reduced from 1.26±0.14 µV at the pre-injection level to 0.66±0.07 µV (P=0.003, t test, 2-tailed), at the same CM level (0.66±0.12 µV) of WT mice (Fig. 4**i**). ABR thresholds in Cx26^+/-^ mice (Fig. 4**j**) but not in WT mice (Fig. 4**k**) were also significantly elevated after PPADS injection. At 8 h after the injection of PPADS, the ABR thresholds in Cx26^+/-^ mice were elevated to the same levels in WT mice and there were no significant differences between Cx26^+/-^ mice and WT mice (P=0.52-0.97, t test, 2-tailed) (Fig. S5). In comparison with WT mice, DPOAE in Cx26^+/-^ mice also had significant reductions after administration of PPADS (Fig. 4**l**).

### Attenuation of hyperacusis by administration of P2x2 antagonist in mice with injection of P2x2 vectors

We also tested the effect of PPADS on hyperacusis generated by overexpression of P2x2 in the cochlea in mice with P2x2 vector injection (Fig. 5). Similar to suppression of hearing oversensitivity observed in Cx26^+/-^ mice (Fig. 4), PPADS also reversibly inhibited the hearing oversensitivity in the mice with P2x2 vector injection (Fig. 5). ASR in the P2x2-vector injection mice was significantly reduced after administration of PPADS (P<0.01, t test, 2-tailed) (Fig. 5**a-c**), while ASR in empty vector injection mice had no significant changes after PPADS injection (P=0.33-0.62, t test, 2-tailed) (Fig. 5**d-e**). At 8 h after injection of PPADS, the peak amplitudes of ASRs in mice with P2x2 vector injection were significantly reduced from 5145.5 mV of pre-injection level to 2745.8 mV (P<0.01, t test, 2-tailed) (Fig. 5**c**), while ASRs in mice with empty vector injection were slightly increased from 1296.8 mV at the pre-injection level to 2822.9 mV (P=0.27, t test, 2-tailed) (Fig. 5**c**). There was a significant difference in ASRs at the pre-injection between P2x2 and empty vector injection mice (P<0.01, t test, 2-tailed) (Fig. 5**c**). However, there was no significant difference in ASRs after PPADS injection between two groups (P=0.84, t test, 2-tailed) (Fig. 5**c**). After 72 h of injection, ASRs in the P2x2 vector injection mice were recovered and returned to the pre-injection level and had significant difference again between two groups (P<0.01, t test, 2-tailed) (Fig. 5**c**).

**Fig. 5.**
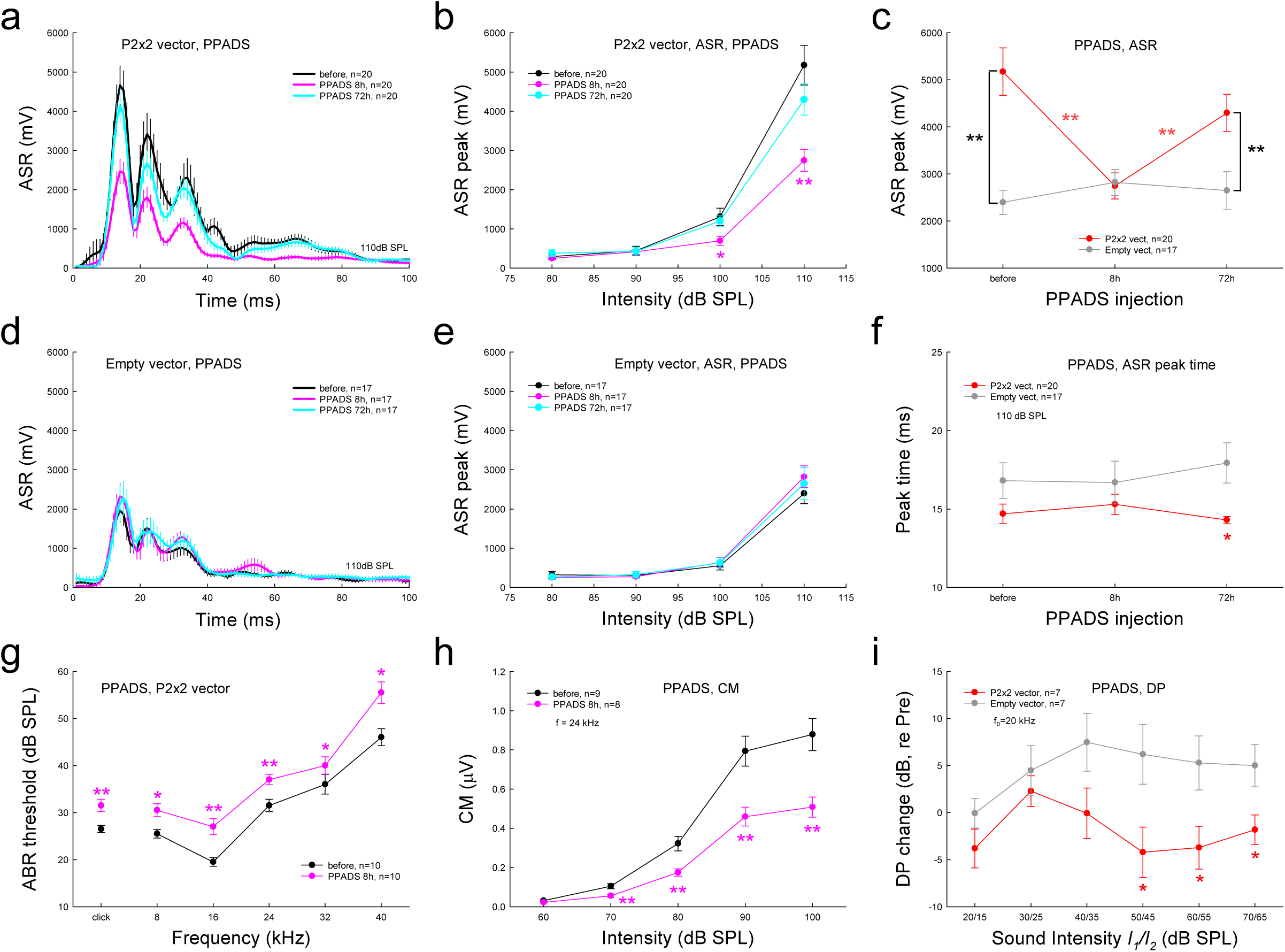
Attenuation of hearing hypersensitivity in P2x2 overexpression mice by administration of PPADS. **a-e**: PPADS reversibly eliminated the enhanced ASR in P2x2 over-expression mice but had no effect on the ASR in WT mice. The ASR in P2x2 overexpression mice was reduced to the WT level after 8 h of administration of PPADS and reversibly recovered at 72 h of PPADS administration (panel **c**). **f**: The effect of PPADS on the ASR peak time. **g-i**: Attenuation of PPADS on hearing hypersensitivity in P2x2 overexpressed ears measured by ABR threshold, CM, and DPOAE. The right ear was injected with P2x2 vectors. After administration of PPADS, the ARB threshold in the right ear was significantly increased and CM and DPOAE in the right ear were significantly reduced. *: P<0.05, **: P<0.01, 2-tail t test.

Also, PPADS had no significant effect on ASR peak time (Fig. 5**f**), except the ASR peak-time in P2x2 vector injection mice at 72 h after injection of PPADS had a significant reduction (P=0.013, t test, 2-tailed) in comparison with empty vector injection mice.

Hearing oversensitivity was also reversibly suppressed by PPADS injection (Fig. 5**g-i**). In the right ear with P2x2 vector injection, ABR thresholds were increased (Fig. 5**g**) and CM and DPOAE were reduced (Fig. 5**h-i**) after administration of PPADS. However, ABR thresholds, CM, and DPOAE in the left ears without vector injection were not significantly changed after administration of PPADS (P=0.1-0.7, t test, 2-tailed) (Fig. S6). Also, at the right ear with empty vector injection in the control mice, ABR thresholds had no significant changes after administration of PPADS (P=0.08-0.76, t test, 2-tailed) (Fig. S7). In comparison with the empty vector injection ears, ABR thresholds in the P2x2 vector injection ears were significantly decreased before injection (Fig. S8**a**). However, after administration of PPADS, there were no significant differences in ABR thresholds between P2x2 and empty vector injection ears (P=0.06-0.76, t test, 2-tailed) (Fig. S8**b**).

### Increase of OHC electromotility in Cx26^+/-^ mice and P2x2 overexpression in the cochlea

The fact of increases of DPOAE in both Cx26^+/-^ mice (Fig. 1**f**) and overexpression of P2x2 in the cochlea (Fig. 2**f**) suggests that active cochlear mechanics are increased. OHCs have electromotility, which is an active cochlear amplification and can increase hearing sensitivity. P2x2 has high expressions at the OHC (Fig. 1**i**). Fig. 6 shows that OHC electromotility associated nonlinear capacitance (NLC) in Cx26^+/-^ mice and P2x2 overexpression mice was increased and left-shifted in comparison with WT mice (Fig. 6**a**). NLC was 6.62±0.18 pF and 6.66±0.24 pF in Cx26^+/-^ mice and P2x2 overexpression, respectively. In comparison with 5.85±0.25 pF in WT mice, they had significantly increased (P=0.02 and 0.02, respectively, one-way ANOVA with a Bonferroni correction) (Fig. 6**e**). Q_max_ in Cx26^+/-^ mice and P2x2 overexpression mice was 0.77±0.01 pC and 0.82±0.04 pC, respectively, and significantly increased as well in comparison with 0.70±0.03 pC in WT mice (P=0.04 and 0.02, respectively, one-way ANOVA with a Bonferroni correction) (Fig. 6**b**). Moreover, the voltage of peak capacitance (V_pk_) in Cx26^+/-^ mice and P2x2 overexpression mice was -74.1±5.47 mV and -82.4±4.31 mV, respectively, showing significant left shift in comparison with -57.5±3.28 mV in WT mice (P=0.02 and 1.58e-4, respectively, one-way ANOVA with a Bonferroni correction) (Fig. 6**d**). However, z was 0.89±0.01, 0.85±0.02, and 0.87±0.02 in Cx26^+/-^ mice, overexpression of P2x2, and WT mice, respectively (Fig. 6**c**), and had no significant changes among three groups (P=0.37-0.41, one-way ANOVA).

**Fig. 6.**
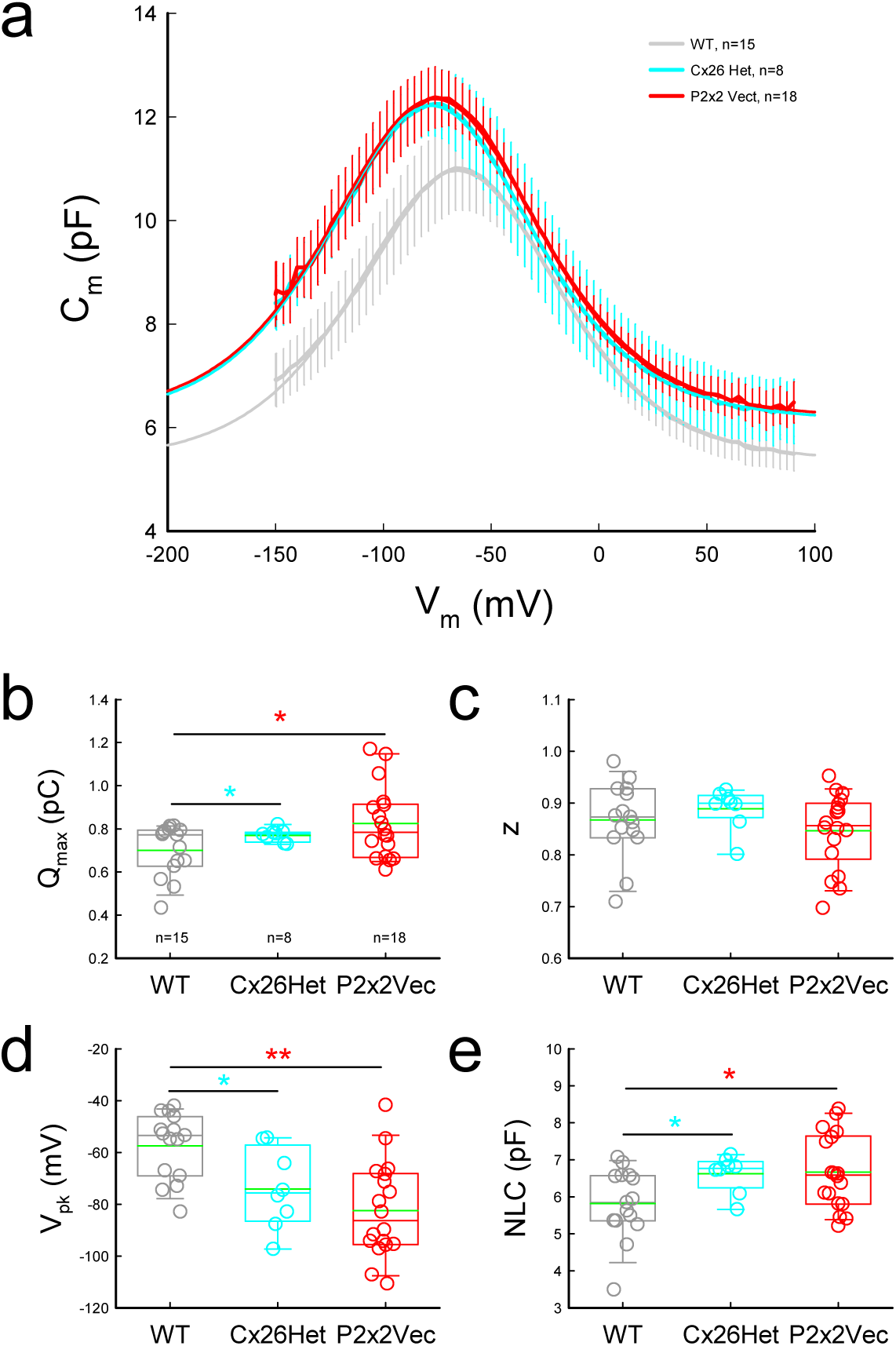
Increase and left-shifting of OHC electromotility associated nonlinear capacitance (NLC) in Cx26^+/-^ mice and P2x2 overexpression in the cochlea by P2x2 vector injection. **a**: Averaged NLCs recorded from WT, Cx26^+/-^ Het, and P2x2 vector injection mice. Erro bars represent SEM. Smooth lines represent fitting by the first derivative of the Boltzmann equation. The fitting parameters in WT, Cx26^+/-^, and P2x2 vector injection mice are: Q_max_=0.70, 0.77, and 0.80 pC; z=0.82, 0.82, and 0.80; V_pk_=-65.7, -77.8, and -76.3 mV; C_lin_=5.36, 6.16, and 6.20 pF, respectively. **b-e**: Parameters of NLC fitting in WT, Cx26^+/-^ Het, and P2x2 vector over-expression mice. In comparison with WT mice, Q_max_ and NLC in Cx26^+/-^ Het and P2x2 vector injection mice were significantly increased and V_pk_ (voltage at peak of NLC) was significantly left-shifted to negative potential. *: P<0.05, **: P<0.01, one-way ANOVA with a Bonferroni correction.

In addition, the expression of prestin in the cochlea had no significant changes in Cx26^+/-^ mice and in P2x2 overexpression mice as well (Fig. S9). Prestin expressions in WT and Cx26^+/-^ mice were 0.0134±0.0025 (n=14) and 0.0143±0.0033 (n=12), respectively. There was no significant difference between them (P=0.74, 2-tail t test, Fig. S9**a**). Prestin was also not upregulated after injection of P2x2 vector to increase P2x2 expression in the cochlea (Fig. S9**a**). The expression of prestin in control mice with empty vectors and P2x2 vector injection mice were 0.0125±0.0028 (n=12) and 0.0148±0.0027 (n=17), respectively (P=0.55, 2-tail t test, Fig. S9**b**).

### Enhancement in mediation of ATP on OHC electromotility by upregulation of P2x2

ATP can mediate OHC electromotility (Yu & Zhao, 2008; Zhao et al., 2005). Consistent with the upregulation of P2x2 in the cochlea in Cx26^+/-^ mice and P2x2 vector injection (Figs. 1**i**&2**c**), the responses of OHC electromotility to ATP in Cx26^+/-^ mice and overexpression of P2x2 in the cochlea were also increased (Fig. 7). Ater application of ATP (50 μM), NLC was reduced and right shifted. In Cx26^+/-^ mice and overexpression of P2x2, the changes of V_pk_ were 7.92±1.38 and 10.8±1.93 mV, respectively, and had significant right-shift in comparison with 7.92±1.38 mV in WT mice (P=0.008 and 0.003, respectively, one-way ANOVA with a Bonferroni correction) (Fig. 7**f**). The NLC was also significantly reduced by -0.62±0.19 and -0.78±0.18 pF, respectively, in comparison with -0.12±0.11 pF in WT mice (P=0.04 and 0.01, respectively, one-way ANOVA with a Bonferroni correction) (Fig. 7**g**). However, there were no significant differences in changes of Q_max_ among three groups (P=0.2-0.8, one-way ANOVA) after application of ATP (Fig. 7**d**). It also had no significant difference in changes of z in Cx26^+/-^ mice (P=0.03, one-way ANOVA with a Bonferroni correction) but not in the P2x2 vector injection group (P=0.24, one-way ANOVA) in comparison with WT mice (Fig. 7**e**).

**Fig. 7.**
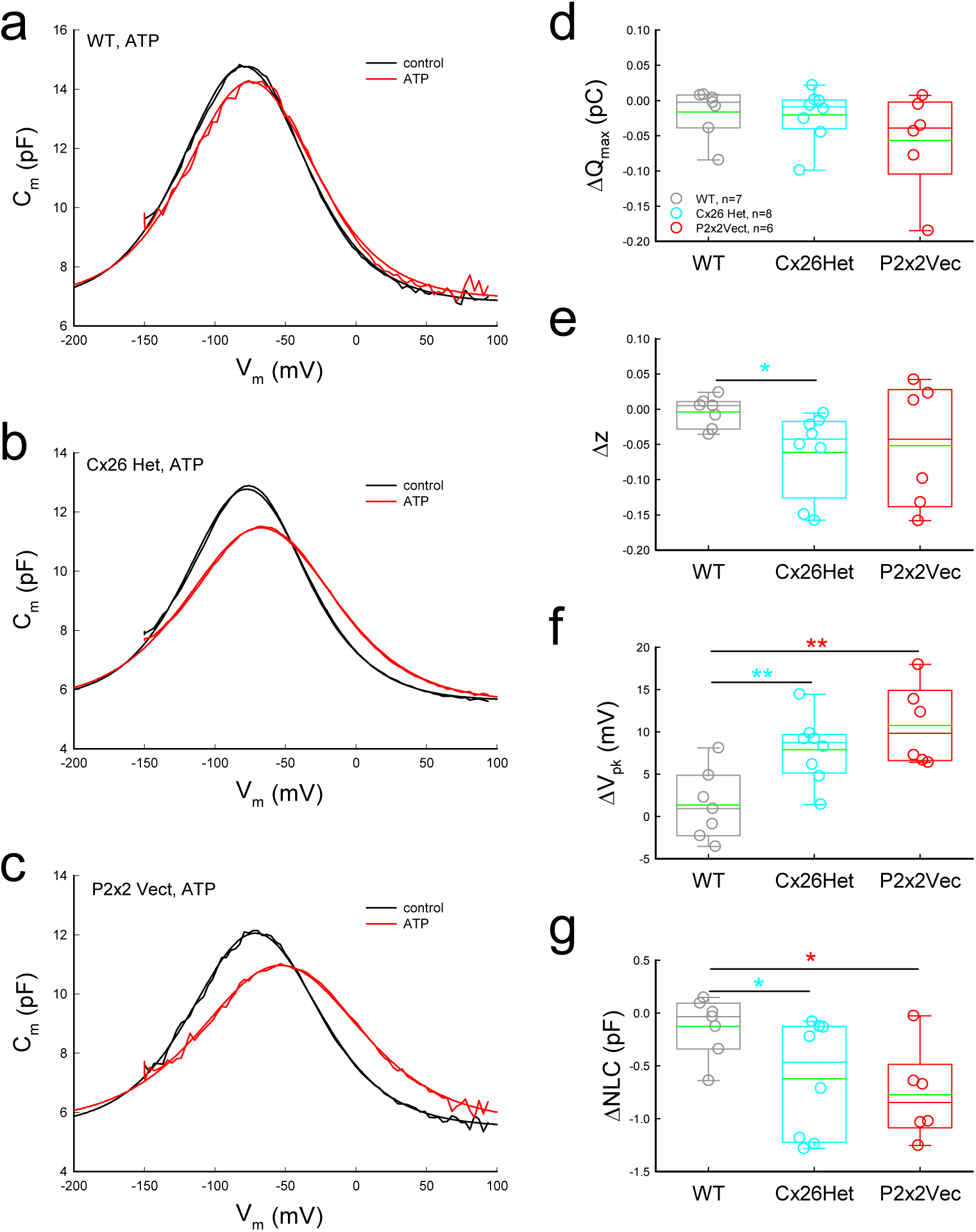
Modification of ATP on OHC electromotility in Cx26^+/-^ mice and P2x2 vector injection mice. **a-c**: NLCs recorded from WT, Cx26^+/-^, and P2x2 vector injection mice at control before ATP application and at 50 μM ATP application. Smooth lines represent fitting with Boltzmann function. The fitting parameters in WT, Cx26^+/-^, and P2x2 vector injection mice are: Q_max_=0.93, 0.82, and 0.80 pC; z=0.88, 0.91, and 0.85; V_pk_= -79.7, - 77.4, and -71.4 mV; C_lin_=6.80, 5.63, and 5.50 pF, respectively, in control, and Q_max_=0.90, 0.82, and 0.79 pC; z=0.85, 0.74, and 0.69; V_pk_= -74.9, -67.6, and -53.4 mV; C_lin_=6.92, 5.57, and 5.67 pF, respectively, after application of 50 μM ATP. **d-g**: Changes in fitting parameters after application of 50 μM ATP. In comparison with WT, ATP-evoked changes of V_pk_ and NLC in Cx26^+/-^ and P2x2 vector injection mice are significantly larger than those in WT mice. *: P<0.05, **: P<0.01, one-way ANOVA with a Bonferroni correction.

P2x2 is also required for medication of ATP on OHC electromotility (Fig. S10). In P2x2 KO mice, the ATP-evoked inward current in the OHC disappeared (Fig. S10**b-c**), and medication of ATP on OHC electromotility was also eliminated (Fig. S10**d-h**). However, different from P2x2 upregulation, downregulation or knockout of P2x2 did not significantly change OHC electromotility associated NLC and ASR (Fig. S11). As shown in Fig. 3 that there were no significant changes in hearing function and ASR in P2x2^+/-^heterozygous KO mice, P2x2 KO mice also had no significant effects on ASR (Fig. S11**a-c**). Similarly, NLC had no significant changes in P2x2^+/-^ heterozygous and P2x2^-/-^KO mice (Fig. S11**d-h**).

### Elimination of enhancement in OHC electromotility and mediation of ATP in Cx26^+/-^mice and P2x2 overexpression mice by PPADS

Fig. 8 shows that after application of PPADS (50 μM), ATP-evoked responses in the OHC were inhibited. The ATP-evoked inward current in the OHC was inhibited by adding PPADS (50 μM) in the application of 50 μM ATP (Fig. 8**a**). The changes in NLC were also inhibited by PPADS (Fig. 8**b-f**). After adding 50 μM PPADS in the application of 50 μM ATP, there were no significant differences in Q_max_, z, V_pk_, and NLC among Cx26^+/-^, P2x2 overexpression, and WT mice (Fig. 8**c-f**).

**Fig. 8.**
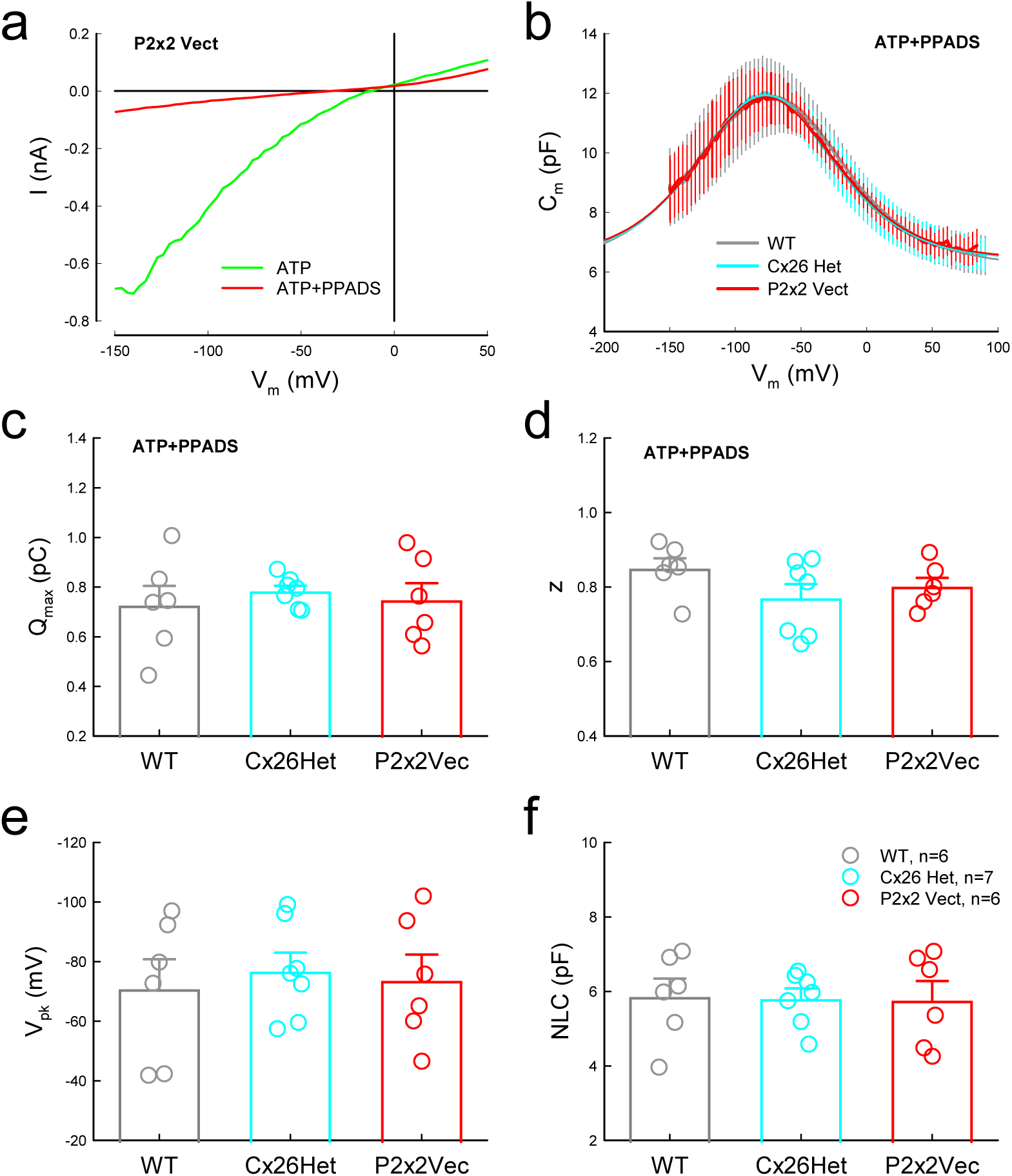
Inhibition by PPADS on enhancement of OHC electromotility in Cx26^+/-^ mice and P2x2 vector injection mice. **a**: ATP-evoked inward current and inhibition by PPADS in an OHC in P2x2 vector injection mice. The OHC was perfused with 50 μM ATP then added with 50 μM PPADS. **b:** Averaged NLC traces recorded from OHCs in WT, Cx26^+/-^, and P2x2 vector injection mice after adding 50 μM PPADS in the application of 50 μM ATP. There are no apparent differences among these recordings. The smooth lines are the Boltzmann functional fitting with parameters: Q_max_=0.85, 0.77, and 0.76 pC; z=0.69, 0.74, and 0.73; V_pk_= -74.6, -76.9, and -76.6 mV; C_lin_=6.22, 6.42, and 6.45 pF in WT, Cx26^+/-^, and P2x2 vector injection mice, respectively. **c-f**: Fitting parameters of Boltzmann function after adding 50 μM PPADS in the application of 50 μM ATP in WT, Cx26^+/-^, and P2x2 vector injection mice. There are no significant differences in fitting parameters among WT, Cx26^+/-^, and P2x2 vector injection mouse groups in one-way ANOVA analysis (P>0.05).

## Discussion

In this study, we found that P2x2 expression in the cochlea but not in the auditory centers was upregulated in hyperacusis generated by Cx26 heterozygous mutations (Figs. 1 and S1&2); overexpression of P2x2 in the cochlea also produced hearing hypersensitivity (Fig. 2). On the other hand, downregulation of P2x2 expression in Cx26^+/-^ mice or administration of P2x2 antagonist PPADS eliminated the hearing hypersensitivity in Cx26^+/-^ mice and P2x2 overexpression mice (Figs. 3-5). OHC electromotility is an active cochlear amplifier and can increase hearing sensitivity. We found that OHC electromotility associated NLC was increased (Fig. 6) and active cochlear mechanics measured by DPOAE (Figs. 1**f**&2**f**) was increased in both Cx26^+/-^mice and P2x2 overexpression mice. The modification of ATP on OHC electromotility in Cx26^+/-^ mice and P2x2 overexpression mice was also enhanced (Fig. 7). Finally, such an increase in OHC electromotility (Fig. 8) and the enhancement in the active cochlear amplification (Fig. 4**l**&5**i**) in Cx26^+/-^ mice and P2x2 overexpression mice were suppressed by administration of PPADS. These data suggest that the activation of P2x2 in the cochlea plays a critical role in the hyperacusis generation. The study also reveals that targeting P2x2 receptors could overcome hyperacusis and provides a potential pharmacological therapy for hyperacusis.

As reported in previous studies (Housley et al., 1999; Yan et al., 2013; Zhao et al., 2005; Zhu & Zhao, 2012), P2x2 has extensive expressions in the cochlea, including expressions in IHCs, OHCs, cochlear supporting cells, and the lateral wall (Fig. 1**i**).

However, no expressions of P2x2 were detectable at the transcriptional level in the AC, IC, and CN by dPCR examinations (Fig. S2**c&d**). P2x2 expressions in the AC, IC, and CN auditory centers also had no significant upregulation in Cx26^+/-^ mice (Fig. S1 and S2**a&b**). These results suggest that upregulation of P2x2 in the cochlea but not in the auditory centers plays a critical role in the hyperacusis generation in Cx26^+/-^ mice.

This concept is further supported by the fact that injection of AAV P2x2 vectors into the cochlea to overexpress P2x2 could generate hyperacusis (Fig. 2). After injection of AAV P2x2 vectors, the expression of P2x2 in the cochlea was increased by 2.75±0.39 folds (Fig. 2**c**), similar to the upregulation of P2x2 in the cochlea of Cx26^+/-^ mice (2.37±0.19 folds) (Fig. 1). Hearing sensitivity and ASR were also significantly increased (Fig. 2**d-i**). However, there was no significant increase in P2x2 expression in the control cochlea with AAV empty vector injection (Fig. 2**c**); the hearing function also had no significant changes (Fig. 2**d-i**). P2x2 vector injection also did not change Cx26 expression in the cochlea; the expression of Cx26 in the right injection cochlea was similar to that in the left control ear without injection (Fig. S4). This result indicates that hyperacusis generation in the P2x2 vector injection ears is not due to alternation of Cx26 expression.

In the cochlea, P2x2 and P2x7 are two major P2x receptors (Housley et al., 2009). Recently, we found that deficiency of P2x7 receptors impaired cochlear efferent function and increased hearing sensitivity and susceptibility to noise (Liang et al., 2025). P2x7 expresses at type II spiral ganglion neurons and synapse areas under IHCs and OHCs in the cochlea (Liang et al., 2025). Deficiency of P2x7 eliminates type II SGN function and reduces cochlear efferent system input, thereby leading to potentiating OHC electromotility and eventually active cochlear amplification (Liang et al., 2025). This is a different pathway from the current study. In this study, we found that there were no changes in P2x7 expression in the Cx26^+/-^ mice (Fig. S3) as well as after injection of P2x2 AAV vectors (Fig. S4), indicating that hyperacusis in Cx26^+/-^ mice is not generated by reduction of P2x7 expression. However, they may share a similar mechanism, i.e., to potentiate OHC electromotility and active cochlear amplification (but with different pathways) increasing hearing sensitivity.

Indeed, we found that the upregulation of P2x2 in the cochlea increased OHC electromotility and enhanced active cochlear amplification (Figs. 1, 2, and 6). Prestin is the OHC electromotility motor protein (Zheng et al., 2000). P2x2 co-expressed with prestin at the OHC lateral wall [Fig. S10**a**, also see (Zhao et al., 2005)]. In both Cx26^+/-^mice and P2x2 over-expression mice, OHC electromotility measured by NLC increased (Fig. 6). The active cochlear amplification measured by DPOAE in Cx26^+/-^ mice (Fig 1**f** & 3**e**) and P2x2 over-expression mice (Fig. 2**f**) were also enhanced. Most importantly, PPADS could inhibit the increase in OHC electromotility (Fig. 8) and the enhancement of DPOAE in both Cx26^+/-^ mice and P2x2 over-expression mice (Figs. 4-5). These data reveal that upregulation of P2x2 in the cochlea to over-increase hearing sensitivity is mainly through increasing OHC electromotility to enhance active cochlear amplification.

In this study, we found that OHC electromotility in Cx26^+/-^ mice and P2x2 overexpression mice was enhanced (Fig. 6). This is consistent with our previous report that prestin expression examined by immunofluorescent staining was increased in Cx26^+/-^mice (Liu et al., 2023). However, dPCR measurement (Fig. S9) showed that prestin expression at the transcriptional mRNA level was not increased. These data indicate that OHC electromotility was enhanced through post-transcription functional modulation.

This concept is further supported by the fact that application of PPADS to inhibit P2x2 receptor function could inhibit increase in OHC electromotility, i.e., the OHC electromotility in Cx26^+/-^ mice and P2x2 overexpression mice had no significant difference from WT mice after administration of PPADS (Fig. 8). The effect of PPADS on OHC electromotility is reversible (Yu & Zhao, 2008). The inhibition of PPADS on hyperacusis was also reversible (Figs. 4&5). Moreover, a recent study demonstrated that OHC electromotility could be functionally modulated by gap junctional coupling between Deiters cells through cochlear efferent system control (Zhao et al., 2022). Thus, OHC electromotility can be mediated by post-transcriptional modulation.

Currently, the mechanism underlying hyperacusis generation remains largely unclear. Little is known about its genetic cues. Our previous study demonstrated that Cx26 heterozygous deficiency in both humans and mouse models can cause hearing oversensitivity (Liu et al., 2023). In this study, we found that hyperacusis in Cx26^+/-^ mice was generated by increasing OHC electromotility and active cochlear amplification through the upregulation of P2x2 expression in the cochlea (Figs. 1&6); overexpression of P2x2 in the cochlea also induced hyperacusis generation by increasing OHC electromotility to enhance active cochlear amplification (Figs. 2&6). These data reveal that P2x2-mediated ATP-purinergic cell signaling mediates OHC electromotility to enhance active cochlear amplification over-amplifying hearing sensitivity, therefore leading to hyperacusis generation. We previously also reported that P2x7 has a critical role in the cochlear efferent system to control hearing sensitivity; P2x7 deficiency can impair type II auditory nerve function and consequently the cochlear efferent inhibition leading to hearing hypersensitivity (Liang et al., 2025). Taken together, these findings suggest that ATP-purinergic cell signaling plays an important role in hearing sensitivity controlling. These findings also provide important information for genetic cues of hyperacusis generation and may be for other related psychological comorbidities as well.

These studies also reveal potential therapy for attenuation of hyperacusis. Currently, no specific pharmacologic treatment for hyperacusis is available in the clinic. In this study, we found that administration of PPADS could inhibit enhancement of OHC electromotility by P2x2 upregulation (Fig. 8) and hearing hypersensitivity in the Cx26^+/-^mice and P2x2 overexpression mice (Figs. 4&5). Downregulation of P2x2 in P2x2^+/-^ /Cx26^+/-^ mice also attenuated hearing hypersensitivity (Fig. 3). Moreover, administration of PPADS had no significant effect on hearing function in WT mice or mice with injection of empty vectors (Figs. 4, 5, and S6-8). These data demonstrate that the ATP-purinergic pathway is a potential and promising target for hyperacusis treatment. This also provides a potential therapeutic strategy for treatment of other psychological disorders, since hearing hypersensitivity can cause many psychological disorders, such as anxiety, learning disabilities, ADHD, post-traumatic stress disorder, and tinnitus (Baguley, 2003; Knipper et al., 2013; Pienkowski et al., 2014; Tyler et al., 2014; Xie et al., 2025).

*GJB2* mutations account for more than 50% of nonsyndromic hearing loss in the clinic. As the most common deafness-associated gene, *GJB2* has a high heterozygous carrier frequency reported to be as high as 20% in the general human populations (Dai et al., 2007; Green et al., 1999; Han et al., 2008; Wattanasirichaigoon et al., 2004; Zelante et al., 1997). We previously reported that both Cx26 heterozygous mutation human carriers and Cx26^+/-^ mice had hearing hypersensitivity (Liu et al., 2023). In this study, we further found that upregulation of P2x2 expression in the cochlea in Cx26^+/-^ mice leads to hearing hypersensitivity by increasing OHC electromotility and active cochlear amplification (Figs. 1&6); the hearing hypersensitivity also could be attenuated by PPADS (Figs. 4&5). These findings are of particular significance in the hyperacusis generation and treatment, given the high prevalence of heterozygous *GJB2* mutations in the general population.

## Materials and Methods

### Animals

All procedures and experiments followed in the use of animals were approved by the University of Kentucky’s Animal Care & Use Committee (UK: 00902M2005) and Yale University’s Institutional Animal Care & Use Committee (Yale: 2022-20463) and conformed to the standards of the NIH Guidelines for the Care and Use of Laboratory Animals. P2x2 knockout (KO) mice were purchased from The Jackson Lab (Stock No: 004603) and were crossed with CBA/CaJ strain (Stock No: 000654, The Jackson Lab) for more than 6 generations. Cx26 deletion was generated by a Cre-FloxP technique. Pax2-Cre mice (the Mutation Mouse Regional Center, Chapel Hill, NC) were crossed with Cx26*^loxP/loxP^* mice (EM00245, European Mouse Mutant Archive) to create Pax2-Cx26 KO mice (Liang et al., 2012). The genotype of mice was identified by tail identification. Both genders of adult mice were used in experiments. WT littermates were used as controls.

Mice were housed in a quiet individual room in basement with regular 12/12 light/dark cycle. The background noise level in the mouse room at mouse hearing range (4-70 kHz) was <30 dB SPL.

### AAV vector injection

The viruses AAV-EFLa-DIO-P2rX2-3FLAG-WPRE (AAV2/9, 1.61 × 10^13^ vg/ml) and its control viruses AAV-EFLa-DIO-mCherry (AAV2/9, 1.47 × 10^13^ vg/ml) were purchased from Obio Technology Corp., Ltd. (Shanghai, China) (Kuang et al., 2022). The AAV vector (0.8 µL) was injected into the inner ear via the posterior semicircular canal (PTSC) by using a micro-injection system (NL2010MC2T, World Precision Instruments, FL, USA).

### Intraperitoneal injection of drug

PPADS (pyridoxalphosphate-6-azophenyl-2’,4’-disulfonic acid) was purchased from Sigma-Aldrich, USA (P178). Before injection, PPADS solution (1 mM) was freshly made and intraperitoneally injected (0.1 mL/10g) into mice.

### RNA-Sequencing and analysis

The auditory cortex, inferior colliculus, cochlear nucleus, and cochlear were quickly isolated and quick-frozen in liquid nitrogen. RNAs were extracted by use of absolute RNA Miniprep Kit (#400800, Agilent, Santa Clara, CA, United States) following manufacture’s instruction. The extracted RNA was examined for quality and concentration by NanoDrop™ Spectrophotometer (ND-ONE-W, Thermo Fisher Scientific Inc.) and sent to Yale Center for Genome Analysis (YCGA) for Bulk Poly(A) RNA sequencing (Zhai et al., 2025).

The raw sequencing reads of RNA-seq experiments were trimmed off sequencing adaptors and low-quality regions by Btrim (Kong, 2011). The trimmed reads were mapped to the mouse genome (GRCm39) by aligner STAR (Dobin et al., 2013) with default parameters. After the counts are collected, the differential expression analysis was done by DEseq2 (Love et al., 2014). The gene symbols were added with the R/Bioconductor package BiomaRt (Durinck et al., 2009). The Kyoto Encyclopedia of Genes and Genomes (KEGG) and Gene Ontology (GO) enrichment analysis was done using R package clusterProfiler (Xu et al., 2024). For AC, IC, and cochlea, there are 10 Cx26 samples and 10 WT samples for each region. For CN, there are 4 Cx26 samples and 4 WT samples.

### Digital droplet PCR

The extracted RNAs (500 ng) were reversely transcribed into DNA by using iScript cDNA Synthesis Kit (Cat. #1708891, Bio-Rad Laboratories). Digital droplet PCR (dPCR) was performed by QuanStudio Absolute Q Digital PCR system (Thermo Fisher Scientific) with Absolute Q DNA digital PCR Master Mix (A52490, Thermo Fisher Scientific) and related mouse gene TaqMan assays (P2x2, Assay ID: Mm00462952_m1; P2x7, Assay ID: Mm001199500_m1; Prestin, Assay ID: Mm00446145_m1, Thermo Fisher Scientific) following instructions. A two-step protocol with 40 cycles was adopted. Mouse TaqMan copy number reference assay Tfrc (4458366, Thermo Fisher Scientific) was employed as an internal reference. Data was analyzed by QuantStudio Absolute Q Digital PCR Software 6.0 (Thermo Fisher Scientific). Changes of gene expressions were normalized to WT mice (Zhai et al., 2025).

### Immunofluorescent staining and confocal microscopy

As described in our previous reports (Liang et al., 2025; Liu et al., 2023; Zhao et al., 2022; Zhao et al., 2021), the mouse cochlea was freshly isolated. The cochlea was incubated with 4% paraformaldehyde in PBS for 0.5-1 h. After fixation, the cochlea was dissected in the normal extracellular solution (NES) (142 NaCl, 5.37 KCl, 1.47 MgCl_2_, 2 CaCl_2_, 10 HEPES in mM, pH 7.4) or embedded in OCT for cross-section. The isolated cochlear sensory epithelia or the cochlear cross-sections were incubated in a blocking solution (10% goat serum and 1% BSA of PBS) with 0.1% Triton X-100 for 30 min.

Then, the epithelia or sections were incubated with primary antibodies for P2x2 (1:250, rabbit anti-P2x2 polyclonal, #P7982, Sigma-Aldrich, USA) or polyclonal goat anti-prestin (1:50, Cat# sc-22694, Santa Cruz Biotech Inc, CA) in the blocking solution at 4°C overnight. After washout, the epithelia were incubated with corresponding second Alexa Fluor antibody (1:400, Thermo Fisher Sci) at room temperature (23 °C) for 2 hr. Before washout, 4’, 6-diamidino-2-phenylindole (DAPI, 50 μg/ml, D1306; Thermo Fisher Sci) was added to visualize the cell nuclei. After incubation for 2 min, the sensory epithelia were washed by PBS 3 times and whole-mounted on the glass-slide. The stained epithelia or slides were observed under a Nikon A1R or AX confocal microscope system (Zhao et al., 2022; Zhao et al., 2021).

### Auditory brainstem response, cochlear microphonic, and distortion product otoacoustic emission recording

Mice were anesthetized by intraperitoneal injection with a mixture of ketamine and xylazine (17.8ml of 0.9% saline +2ml of ketamine + 0.2 ml of xylazine). Body temperature was maintained at 37–38C by placing anesthetized mice on an isothermal pad. Auditory brainstem response (ABR) was recorded in a double-wall sound-isolated chamber by use of a Tucker-Davis R3 workstation (Tucker-Davis Tech. Alachua, FL) (Liu et al., 2023; Mei et al., 2017; Zhao et al., 2022; Zhu et al., 2015; Zhu et al., 2013). High frequency speakers ES-1 and EC-1 (Tucker-Davis Tech. Alachua, FL) were used in open-field recording and close-field recording, respectively. ABR was recorded by stimulation with clicks in alternative polarity and tone bursts (4-40 kHz) from 80 to 10 dB SPL in a 5 dB step in a double-walled sound isolation room. The signal was amplified (50,000x), filtered (300-3,000 Hz), and averaged 500 times. The ABR threshold was determined by the lowest level at which an ABR can be recognized.

CM was evoked by 8-24 kHz tone bursts and recorded with the same electrode setting as ABR recording as previously reported (Liu et al., 2023; Zhu et al., 2013). The signal was amplified by 50,000 with 3-50 kHz filter and averaged by 100 times.

Distortion product otoacoustic emission (DPOAE) was recorded with EC-1 high frequency speaker by using a Tucker-Davis R3 workstation (Tucker-Davis Tech.

Alachua, FL) (Liu et al., 2023; Mei et al., 2017; Zhu et al., 2015; Zhu et al., 2013). Two pure tones (f_1_ and f_2_, f_2_/f_1_=1.22) were simultaneously delivered into the external ear canal through two plastic tubes and sealed with an earplug. The frequencies of two-testing sounds were determined by a geometric mean of f_1_ and f_2_ (f_0_ = (f_1_ × f_2_)^1/2^) at f_0_ = 4, 8, 16, and 20 kHz with f_2_/f_1_ = 1.22. The intensity of f_1_ (I_1_) was set at 5 dB SPL higher than that of f_2_ (I_2_). The distortion product was recorded with an average of 150 times and a cubic distortion product of 2f_1_ – f_2_ was measured as DPOAE.

### Acoustic startle response recording

Acoustic startle response (ASR) was recorded with a computer-controlled SR-LAB Startle Response System (San Diego Instruments, San Diego, CA). Before the test, the mouse was placed in a restraint in the test box in quiet for 2 minutes. ASR was evoked by a series of white-noise pulses (25-ms duration) from 80 to 110 dB SPL in a 10 dB step (Liang et al., 2025; Zhao et al., 2025).

### Patch-clamp recording and nonlinear capacitance measurement

OHCs were freshly isolated from the mouse cochlea (Zhu et al., 2013). The classical whole-cell recording was performed. A patch pipette was filled with an intracellular solution (140 KCl, 10 EGTA, 2 MgCl2, 10 HEPES in mM; 300 mOsm and pH 7.2) and had an initial resistance of 2.5–3.5 MΩ in the bath solution (130 NaCl, 5.37 KCl, 1.47 MgCl_2_, 2 CaCl_2_, and 10 HEPES in mM; 300 mOsm and pH 7.2) (Yu and Zhao, 2008). The patch clamp recording was performed under a whole-cell configuration using Axopatch 200B patch clamp amplifier (Molecular Devices, CA) with jClamp (Scisft, New Haven, CT). Nonlinear capacitance (NLC) was measured by a two-sinusoidal method and fitted to the first derivative of a two-state Boltzmann function with jClamp and MATLAB (Yu & Zhao, 2008; Zhu et al., 2013).

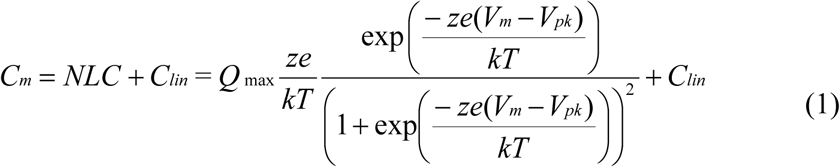

where *Q_max_* is the maximum charge transferred, *V_pk_* is the peak of NLC, *z* is the number of elementary charge (*e*), *k* is Boltzmann’s constant, and *T* is the absolute temperature. Membrane potential (*V_m_*) was corrected for electrode access resistance (*R_s_*).

### Reproducibility, data processing, and statistical analysis

The number of recording mice in each experiment was indicated in the related figure. Each experiment was repeated at least three times. Data were plotted by SigmaPlot and statistically analyzed by SPSS v25.0 (SPSS Inc., Chicago, IL, United States). Data were expressed as mean ± SEM. Parametric and nonparametric data comparisons were performed using one-way ANOVA or Student t tests after assessment of normality and variance (SPSS, SPSS Inc). The threshold for significance was α = 0.05. ANOVAs used Bonferroni *post hoc* test.

## Acknowledgement

This work was supported by NIH Grants R01 DC017025 and R01 DC019687 to HBZ.

## Author contributions

HBZ designed experiments. TYZ, CL, JC, JY, YK, YZ, NY, and HBZ performed experiments. TYZ, CL, JC, JY, YK, and HBZ analyzed data. TYZ and HBZ wrote paper. CL, JC, JY, YK, YZ, and NY reviewed the paper.

## Data and materials availability

All data needed to evaluate the conclusions in the paper are present in the paper and/or the Supplementary Materials.

---

**Fig. S1.**
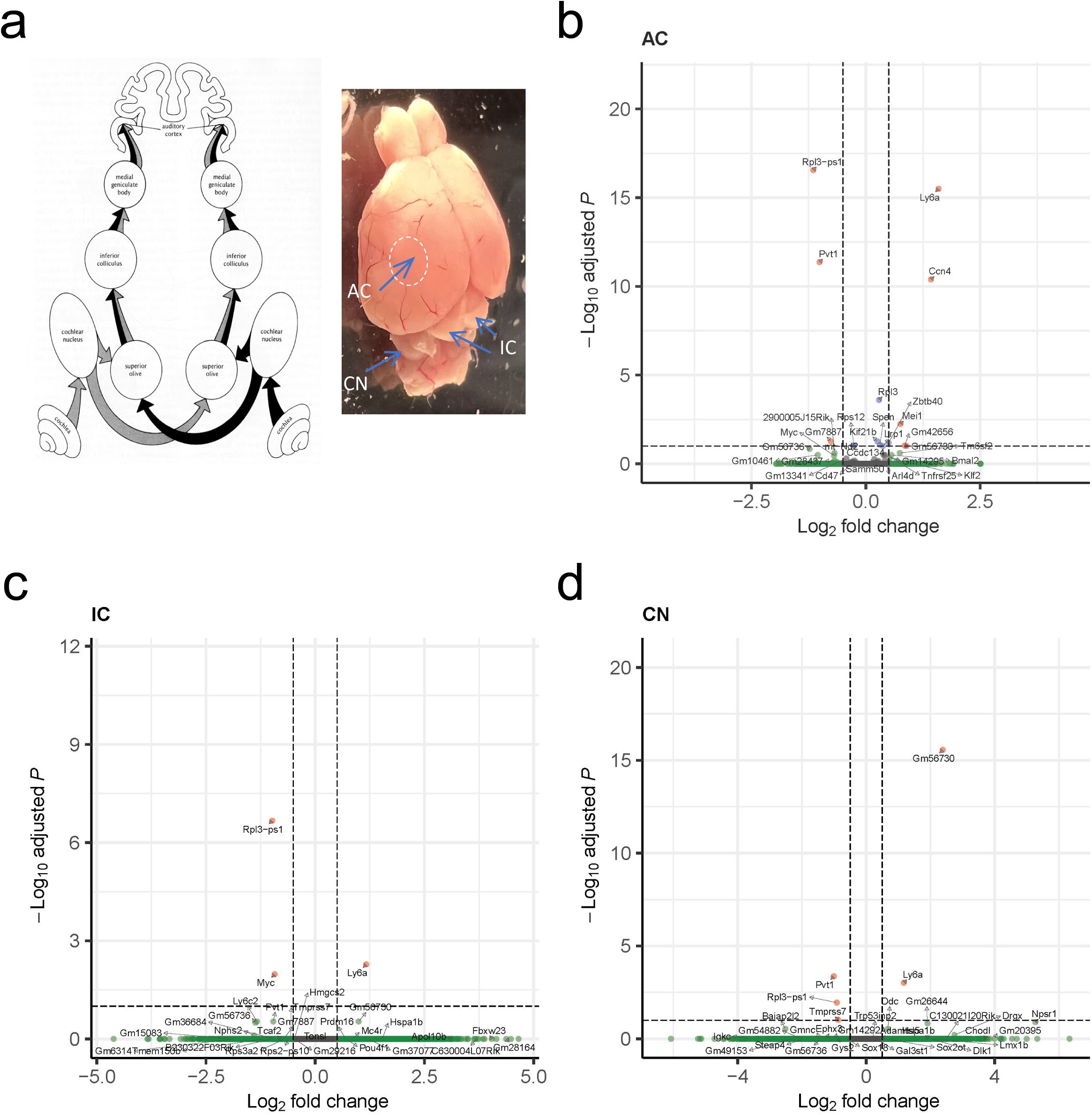
No apparent changes of P2x2 expressions in the auditory centers. **a**: A diagram of the afferent auditory system and locations of the auditory cortex (AC), inferior colliculus (IC), and cochlear nucleus (CN) in the mouse brain (the cerebellum was removed to visualize the CN). **b-d**: Volcano plots of gene upregulation and downregulation in the AC, IC, and CN in Cx26^+/-^ mice in Bulk Poly(A) RNA-Seq. There were no significant changes of P2x2 expressions in these auditory centers in Cx26^+/-^ mice.

**Fig. S2.**
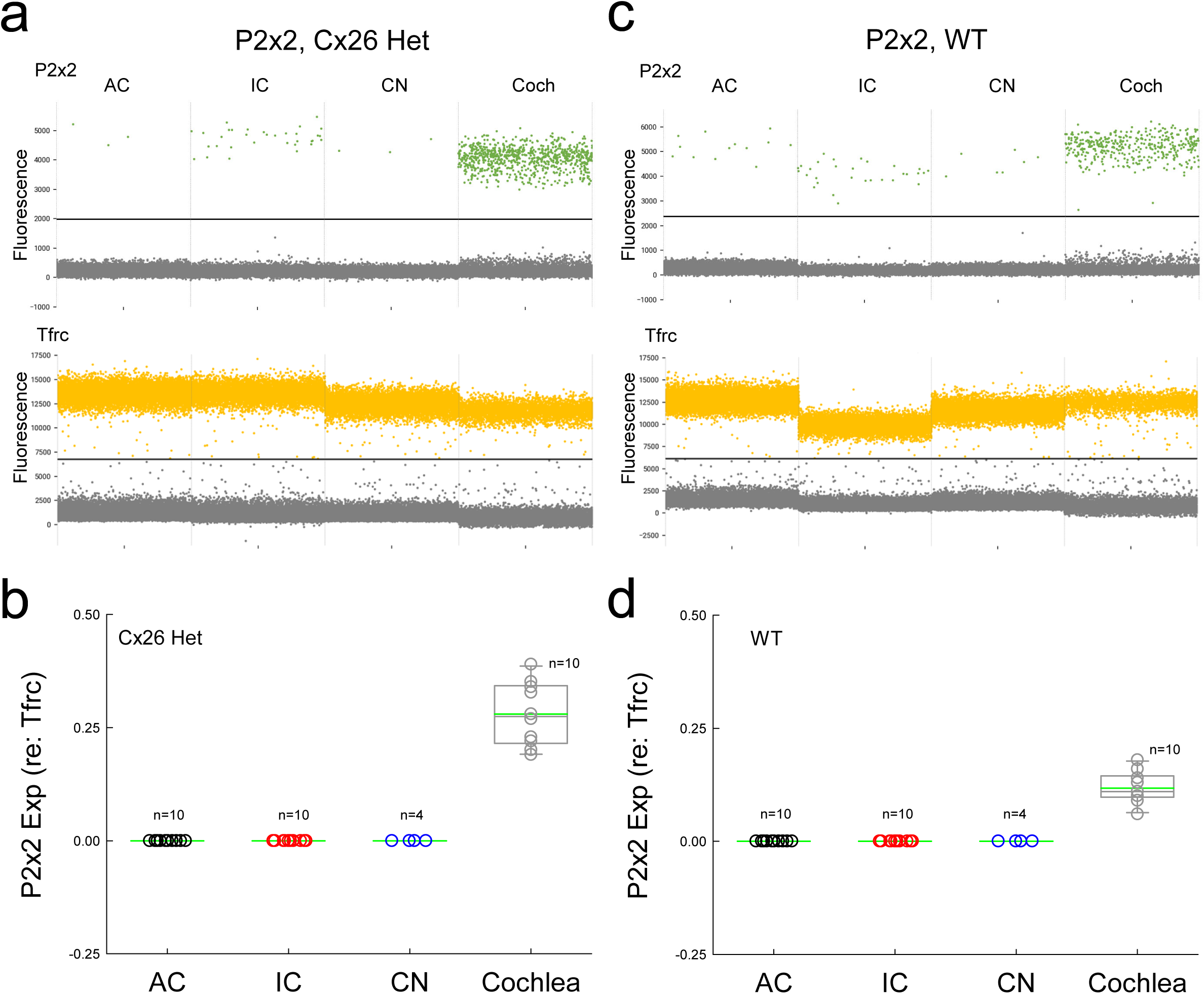
No apparent expression or upregulation of P2x2 expression in the auditory centers in WT and Cx26^+/-^ mice examined by dPCR. The P2x2 expression was calculated from reference to the reference gene *Tfrc*. **a-b**: P2x2 expressions in AC, IC, CN, and cochlea in Cx26^+/-^ mice. **c-d**: P2x2 expressions in the AC, IC, CN, and cochlea in WT mice. Panel **a**&**c** are the represented droplet plots of dPCR for P2x2 and the reference gene *Tfrc* in WT and Cx26^+/-^ mice. There are no P2x2 expressions in the AC, IC, and CN in both Cx26^+/-^ mice and WT mice. In the cochlea, P2x2 expressions referring to Tfrc are 0.28±0.02 and 0.12±0.01 in Cx26^+/-^ mice and WT mice, respectively. In comparison with WT mice, the P2x2 expression in the Cx26^+/-^ mouse cochlea has a significant increase (P>0.001, 2-tail t test).

**Fig. S3.**
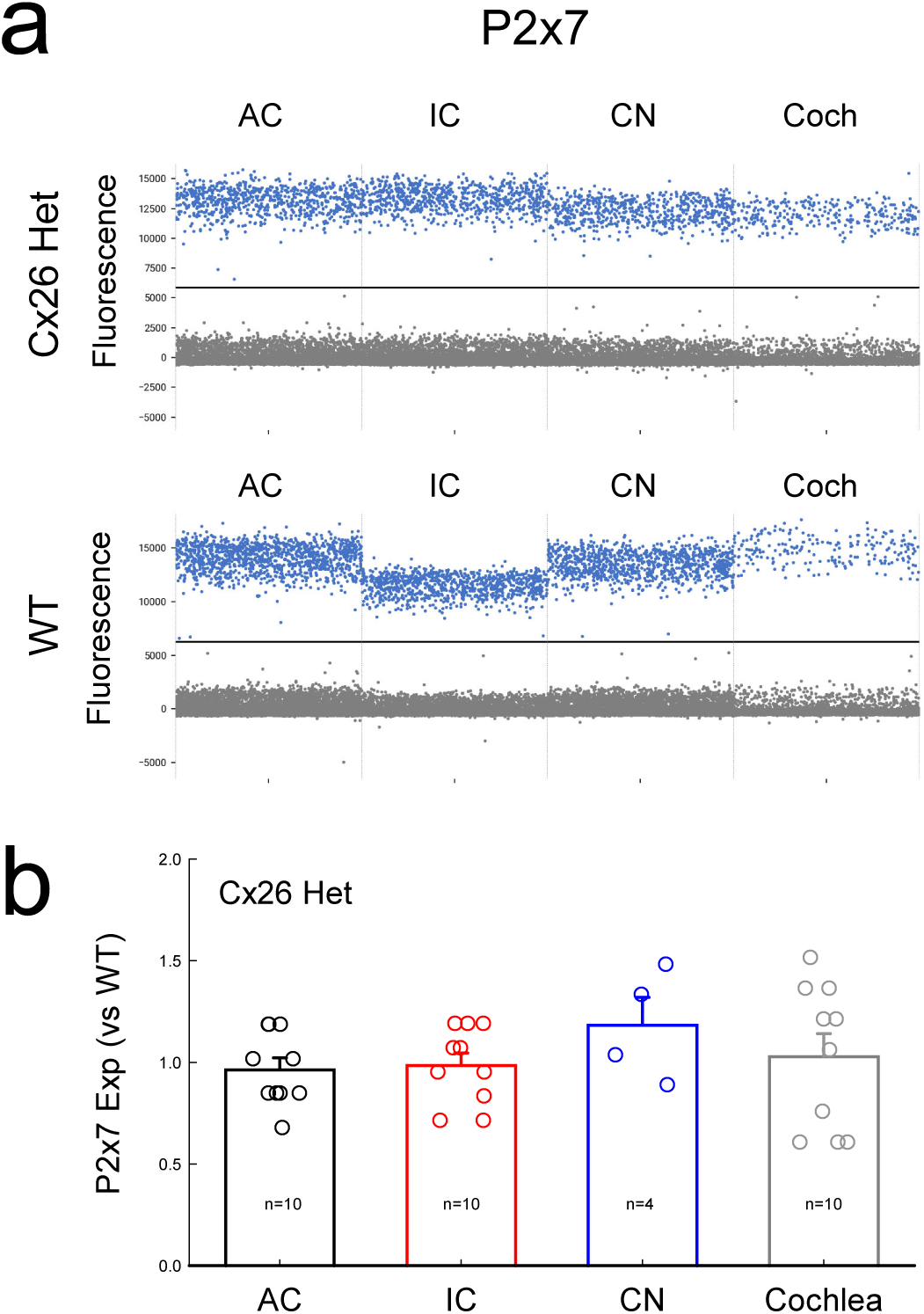
No changes of P2x7 expression in the auditory system in Cx26^+/-^ mice. The expressions of P2x7 in the AC, IC, CN, and cochlea were examined by dPCR. **a**: Represented dPCR droplet plots of P2x7 expressions in the AC, IC, CN, and cochlea in Cx26^+/-^ mice and WT mice. **b**: Fold changes of P2x7 expression in Cx26^+/-^ mice. Referring to WT mice, there are no significant changes of P2x7 expression in Cx26^+/-^mice (P>0.05, one-way ANOVA).

**Fig. S4.**
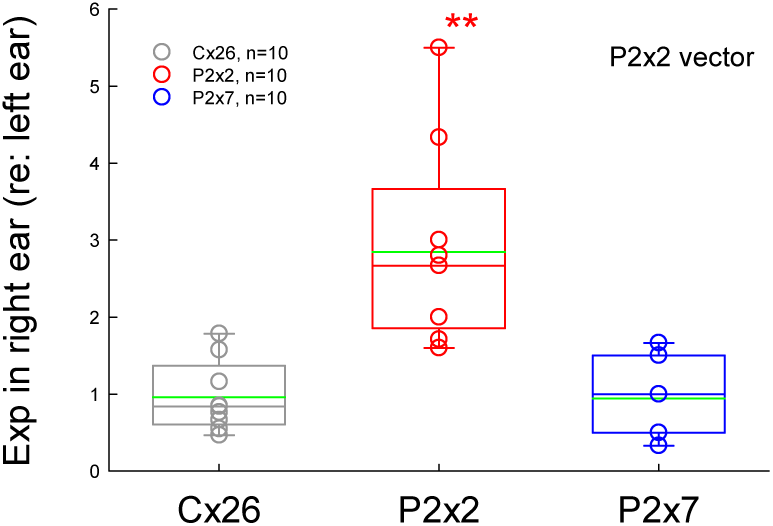
No changes of Cx26 and P2x7 expressions in the cochlea after injection of P2x2 vectors. The expressions of Cx26, P2x2, and P2x7 in the right ears with P2x2 vector injection were measured by dPCR and normalized to those in the left ears without injection. P2x2 expression has a significant increase of 2.85±0.43 (n=10) referring to the referring to the left ears (P<0.01 2-tail t test), while the expressions of Cx26 and P2x7 are 0.94±0.17 and 0.96±0.15, respectively (P=0.75 and 0.81, respectively, 2-tail t test) and have no significant changes.

**Fig. S5.**
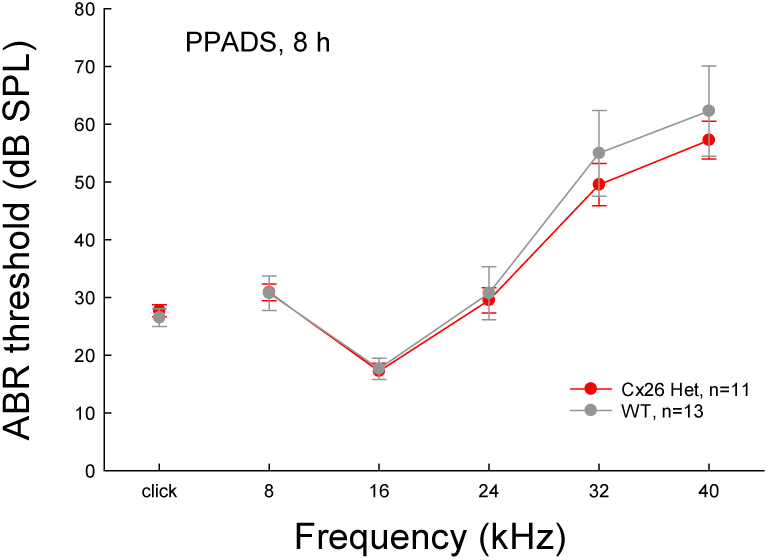
No significant difference in ABR thresholds between Cx26^+/-^ mice and WT mice after administration of PPADS. ABR was measured at 8 h after administration of PPADS. There were no significant differences in ABR thresholds between Cx26^+/-^ mice and WT mice (P = 0.52 – 0.97, 2-tail t test).

**Fig. S6.**
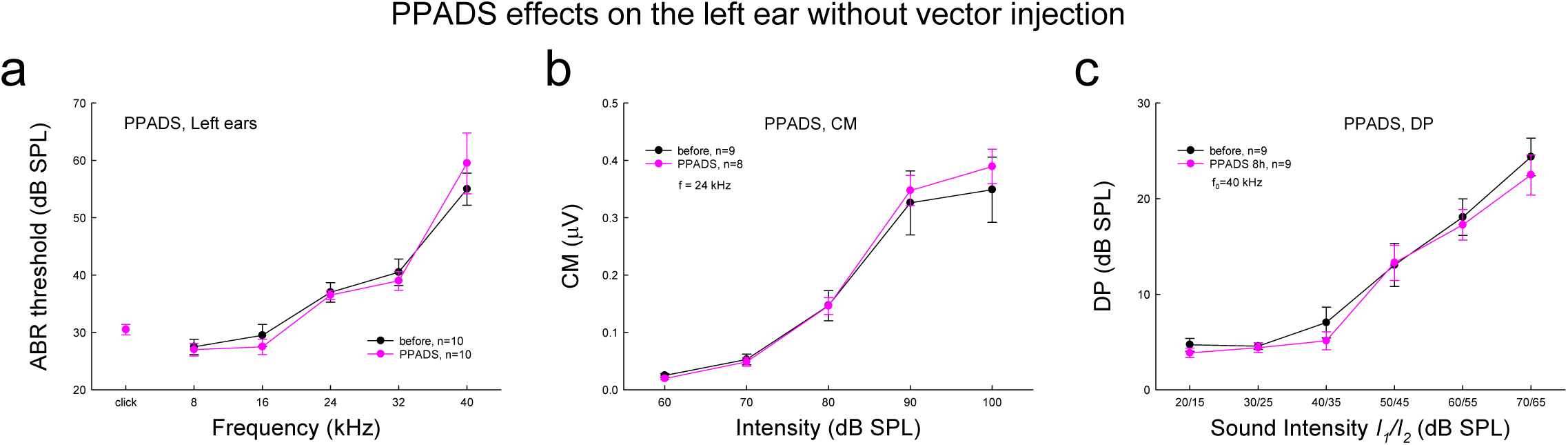
No significant effects on hearing function in the left control ear without P2x2 vector injection after administration of PPADS. Hearing functions in the left control ears were measured before and after 8 h of administration of PPADS by close-field recording. **a-c**: There were no significant changes in ABR, CM, and DPOAE in the left-ears without injection after administration of PPADS.

**Fig. S7.**
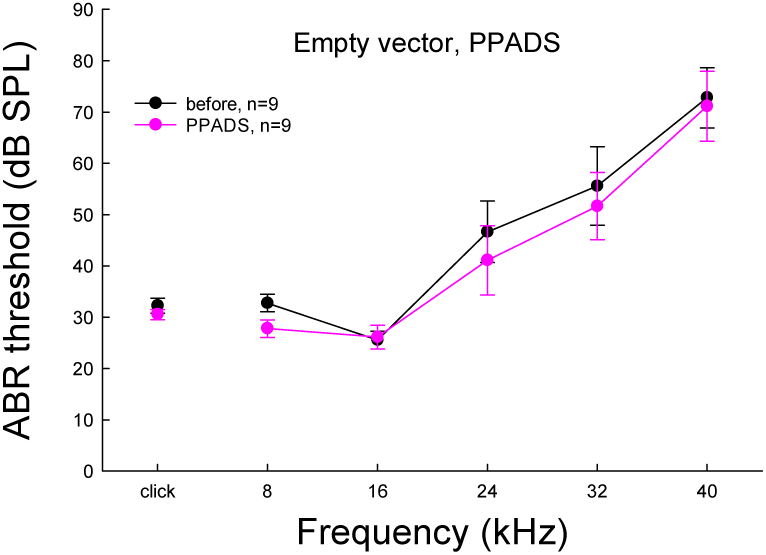
PPADS has no significant effects on hearing function in the right ear with empty vector injection. Right ears were injected with AAV empty vectors without P2x2. ABR thresholds were measured with the close-field recording before and after 8 h of administration of PPADS.

**Fig. S8.**
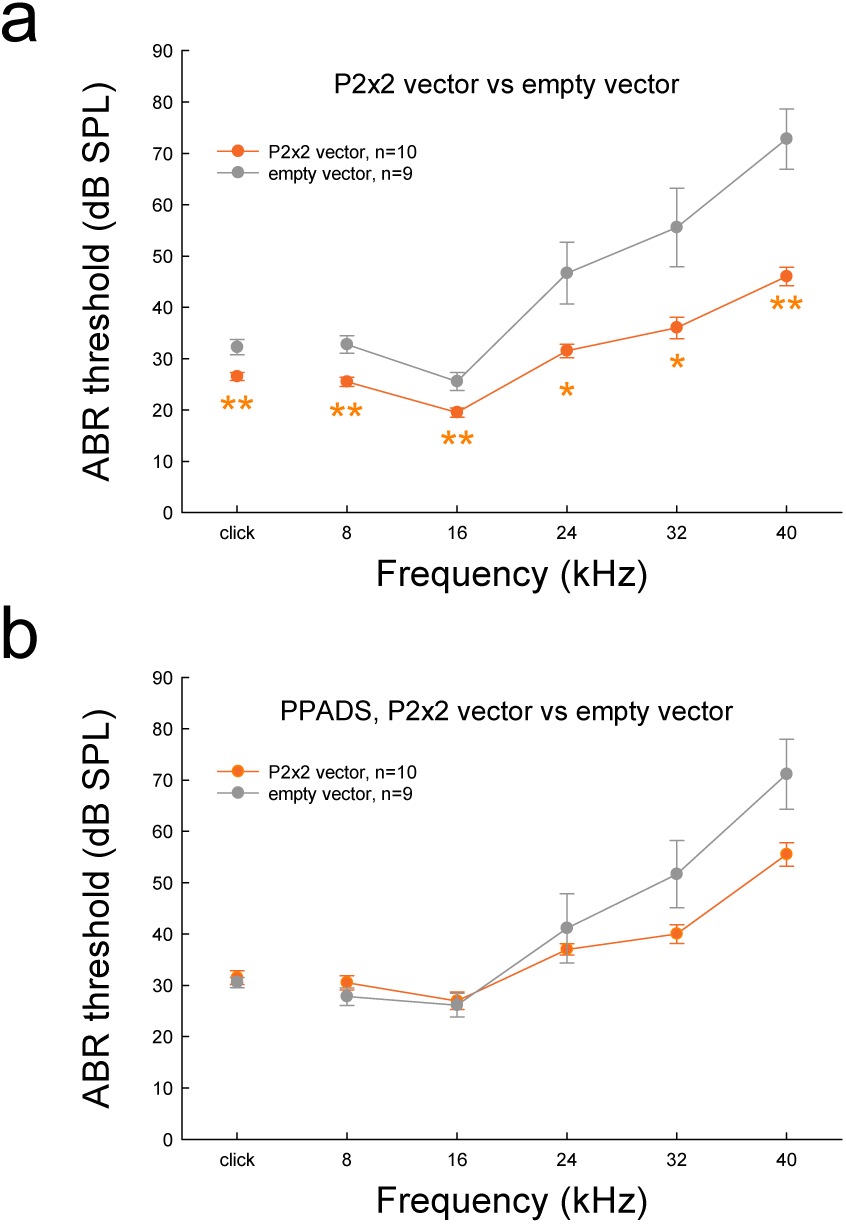
No significant differences of ABR thresholds between mice with P2x2 and empty vector injections after administration of PPADS. The P2x2 or empty vector was injected into only the right ear. ABR thresholds in the right ear were recorded before and after 8 h of administration of PPADS by close-field recording. **a**: ABR thresholds in the right ear with P2x2 or empty vector injection. In comparison with empty vector injections, the injection of P2x2 vectors significantly reduces ABR thresholds. *: P<0.05, **: P<0.01, 2-tail t test. **b**: There are no significant differences in ABR thresholds between P2x2 and empty vector injection ears after 8 h of administration of PPADS.

**Fig. S9.**
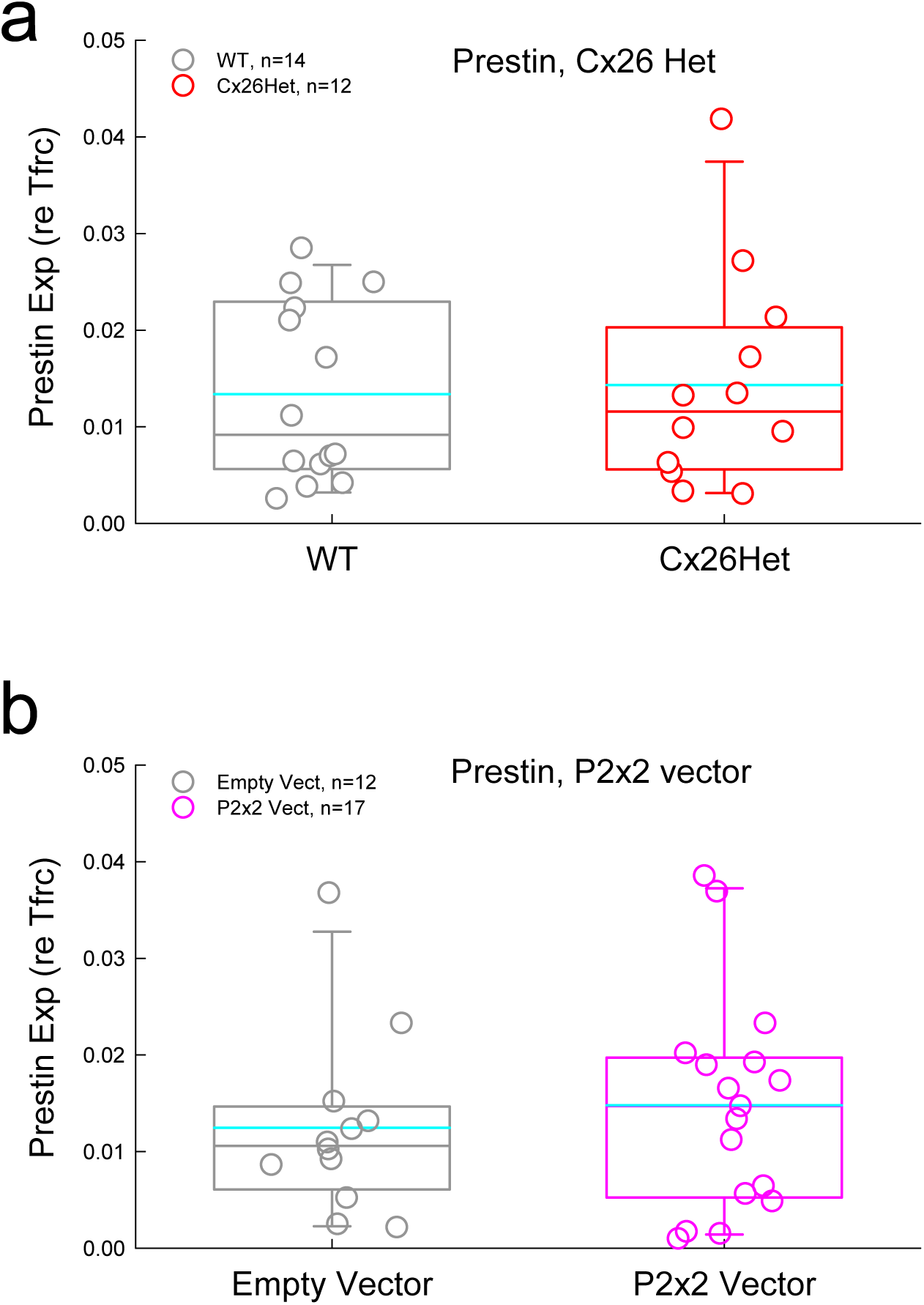
No significant changes in prestin expression in Cx26 hetero-deletion mice and P2x2 vector injection mice. Prestin expression in the cochlea was examined by dPCR and normalized to the reference gene *Tfrc*. **a**: There are no significant changes of prestin in Cx26 hetero-deletion mice. **b**: There are no significant changes of prestin after injection of P2x2 vectors.

**Fig. S10.**
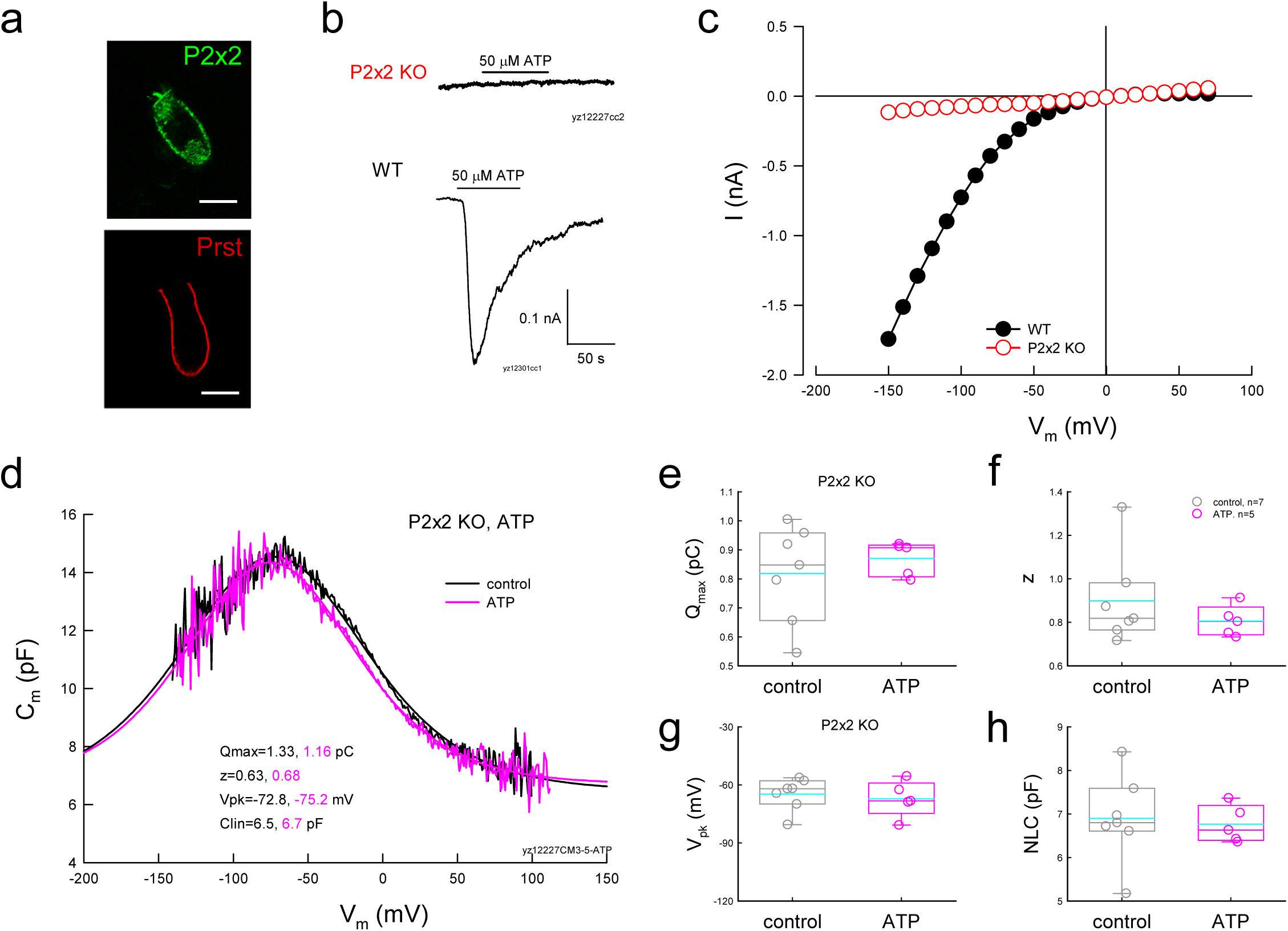
P2x2 receptors are required for ATP modification on OHC electromotility. **a**: Immunofluorescent staining for P2x2 and Prestin in OHCs. Scale bar: 10 μm. **b-c**: ATP-evoked inward current in OHCs and absence in P2x2 KO mice. The OHC was held at -80 mV under the whole-cell recording configuration in patch clamp recording. Application of 50 μM ATP evoked an apparent inward current in OHC in WT mice but not in P2x2 KO mice. There was also no apparent current visible in I-V curve for voltage-step stimulation (panel **c**). **d-h**: There is no apparent effect of ATP on OHC electromotility in P2x2 KO mice. NLCs have no significant changes after perfusion of 50 μM ATP in a P2x2 KO mouse OHC. In comparison with control before perfusion, Q_max_, z, V_pk_, and NLC have no significant changes after perfusion of 50 μM ATP (P>0.05, 2-tail t test).

**Fig. S11.**
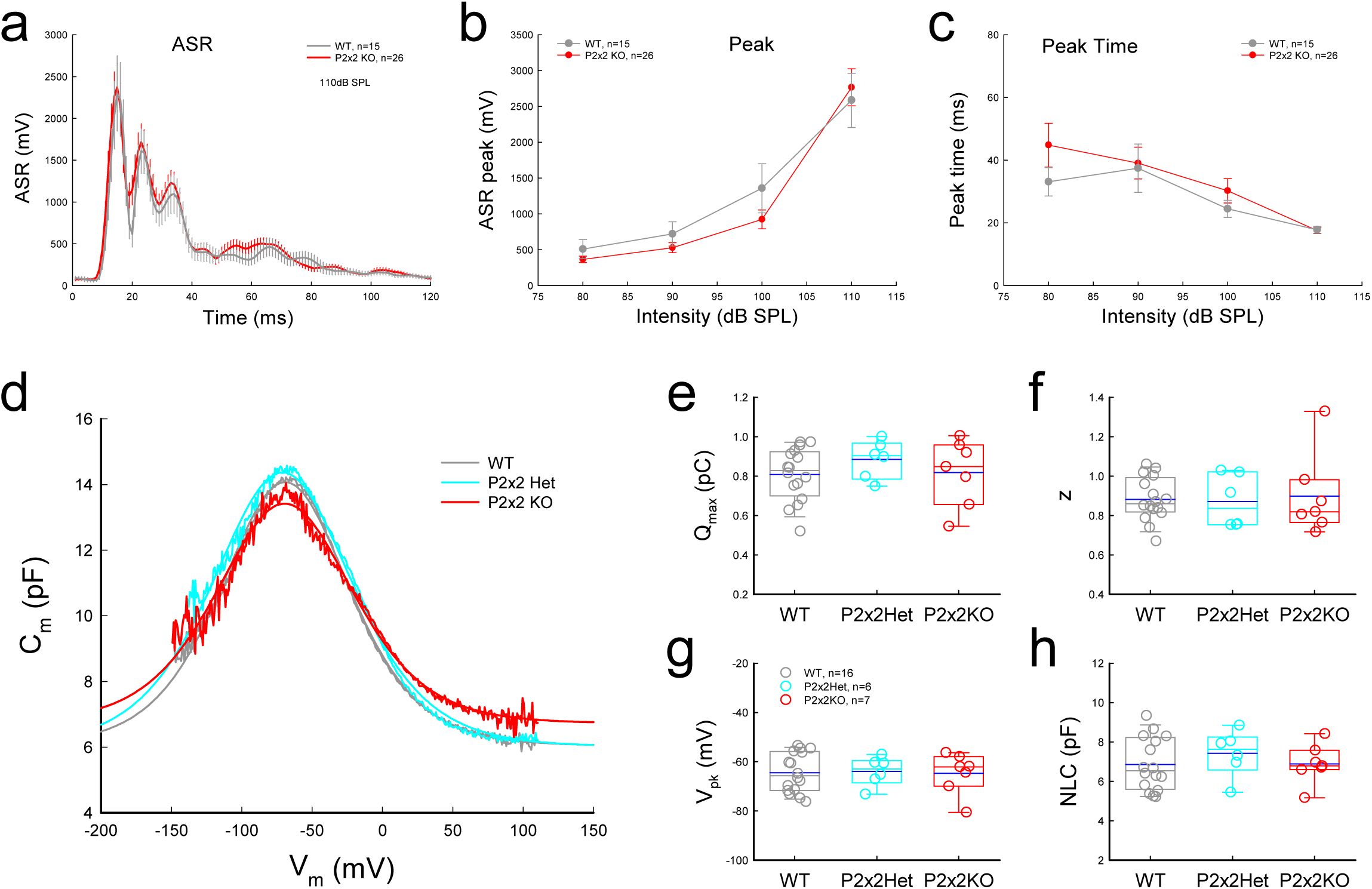
No significant changes of ASR and OHC electromotility in P2x2 deficient mice. **a-c**: No significant changes of ASR in P2x2 KO mice. **d:** NLCs recorded from WT, P2x2^+/-^ hetero-deletion, and P2x2^-/-^ KO mice. Smooth lines represent fitting the first derivative of the Boltzmann equation. The fitting parameters are: Q_max_=0.85 and 0.86 pC; z=0.85and 0.83; V_pk_=-78.0 and -74.6 mV; C_lin_=5.47 and 5.53 pF, respectively, for NLC recorded from WT, P2x2^+/-^, and P2x2 KO mice. **e-h**: Parameters of NLC fitting in the WT, P2x2^+/-^, and P2x2 KO mice. There are no significant changes in OHC electromotility in P2x2^+/-^ and P2x2 KO mice with one-way ANOVA analysis.

